# Ni(II) binding affinity and specificity of solute binding proteins: the importance of the double His motif and variable loop revealed by structural and mutational studies

**DOI:** 10.1101/2022.10.03.510666

**Authors:** Patrick Diep, Peter J. Stogios, Elena Evdokimova, Alexei Savchenko, Radhakrishnan Mahadevan, Alexander F. Yakunin

## Abstract

Extracytoplasmic solute binding proteins (SBPs) are molecular shuttles involved in the cellular uptake of various small molecules and metal ions including Ni(II). Our previous study with the Ni(II) binding proteins (NiBPs) *Cj*NikZ from *Campylobacter jejuni* and *Cc*NikZ-II from *Clostridium carboxidivorans* demonstrated they were able to bind Ni(II) at low micromolar affinity without the need for additional chelators. Here, we determined the crystal structure of apo *Cc*NikZ-II, which revealed a Ni(II) binding site comprised of the highly conserved double His (HH-)prong (His511, His512) and a short variable (v-)loop nearby (Thr59-Thr64, TEDKYT). Alanine scanning mutagenesis of the *Cc*NikZ-II Ni(II) binding site identified Glu60 and His511 as essential for high affinity binding to Ni(II). Phylogenetic analysis of >4000 SBP sequences demonstrated the presence of two clusters of proteins containing the HH-prong with *Cc*NikZ-II and *Cj*NikZ. To provide insights into the role of the double His-prong and v-loop sequence in Ni(II) binding of NiBPs, nine purified *Cc*NikZ-II homologues containing the HH-prong and v-loop were screened using an automated screening workflow. Metal binding assays with purified homologous NiBPs revealed high Ni(II) binding affinity without requirement for chelators indicating that the double His prong represents a signature motif for the presence of Ni(II) binding activity in SBPs. The engineered *Cc*NikZ-II variants with the wild type v-loop (TEDKYT) replaced with v-loops from NiBPs with higher affinity showed up to an order of magnitude higher affinity for Ni(II). In addition, the v-loop appears to play a role in metal ion specificity as purified wild type and engineered NiBPs with different v-loop sequences showed distinct metal profiles. This work paves way for metalloprotein engineering of NiBPs towards biocatalytic and metal recovery applications.

## Introduction

Metal ions (metals) are an indispensable part of all cells due to their unique properties that enable more complex biochemistry. In particular, several transition metals of the first-row d-block elements like Cu(II), Fe(II, III), and Ni(II) (nickel) are indispensable co-factors for the activity of many enzymes. These metals are thus critical for bacterial adaptation to challenging environments such as pathogenesis in the anaerobic gastrointestinal tract of humans, resistance to extremely acidic mine waters, and survival in metal-scarce soil microenvironments (1–3). Bacteria have evolved elaborate and specific mechanisms to bind metals and control their metal homeostasis (*i.e*., detection by metal sensors, active transport across membranes, and trafficking via metallochaperones) (4–7). The periplasmic and extracellular solute binding proteins (SBPs) of ATP binding cassette (ABC) transporters specifically function as molecular tools for binding specific substrates (like metals) from the environment. There are >750 three-dimensional structures of unique SBPs in the Protein Database (PDB) as of July 2022. Specifically, subcluster C-I is comprised of 14 metal-specific SBP structures (8–10). This structural approach (Berntsson-Poolman Classification) to binning SBPs into groups is complemented by the phylogenetic approach (Tam-Saier Classification) to binning these SBPs. Immense diversity (>147K members) can be observed among Family 5, which is comprised of peptide and nickel-specific SBPs, herein termed PepBPs and NiBPs, respectively (11).

Several NiBPs have already been biochemically and structurally characterized. *Ec*NikA from *Escherichia coli* is the canonical NiBP belonging to the most studied metal-specific ABC transporter, NikABCDE (12–19). Several homologous proteins from pathogenic bacteria have also been studied, including *Cj*NikZ from *Campylobacter jejuni* (20, 21), *Bs*NikA from *Brucella suis* (21), *Yp*YntA from *Yersinia pestis* (21), *Sa*NikA and *Sa*CntA from *Staphylococcus aureus* (22), *Hh*NikA from *Helicobacter hepaticus* (23), *Hp*CeuE from *Helicobacter pylori* (24), *Vp*NikA from *Vibrio parahaemolyticus* (25), and *Cd*OppA from *Clostridium difficile* (26). Two common threads tied these studies together: the methodology to determine these NiBPs’ affinity for nickel, and the use of crystal structures to draw conclusions about the NiBPs’ binding mechanisms for nickel. Studies using isothermal titration calorimetry (ITC) (14, 21) and cuvette-based spectrofluorometry of intrinsic tryptophan fluorescence quenching (ITFQ) have confirmed that NiBPs have low micromolar affinities (apparent *K*_D_ ≤ 20 μM) for nickel (13, 18, 20, 24). Binding affinity data combined with crystal structures of NiBPs have painted informative narratives about the binding mechanism of NiBPs, which is necessary for studying microbial pathogenesis.

However, a different approach is needed to engineer NiBPs and metalloproteins in general to have desirable industrial properties: they should possess tolerance to harsh temperatures, pH, and salinity; robust structural integrity to enable multiple re-use cycles; and tuneable affinity and selectivity. Metalloproteins involved in naturally-occurring metal homeostasis systems have been subjects of engineering endeavours for metal removal and recovery applications (27–31). Synthetic biology undertakings are reliant on iterative design cycles and are thus contingent on rapid automation-enabled high-throughput characterization, or at least the development of methods that leverage features of the biological entity/system that simplify screening workflows (*e.g*., spectrofluorometric changes in response to metal titration) (32, 33). In prior work, we explored the scalable use of microplate-based ITFQ by refining protocols based on existing literature (34–37). We first accurately determined the apparent *K*_D_ value of *Cj*NikZ for nickel as a positive control, then determined *K*_D_ values for nickel of *Cc*NikZ-II from *Clostridium carboxidivorans*, which is the second NiBP in the *C. carboxidivorans* acetogenesis operon (37). We found that *Cc*NikZ-II could bind nickel with a *K*_D_ of 13.4 μM (95% CI: 10.7 – 16.1 μM), which is close to that of *Cj*NikZ (*K*_D_ 3.3 μM, 95% CI: 2.4 – 4.3 μM).

In our recent work, we demonstrated that *Cc*NikZ-II and *Cj*NikZ exhibited micromolar affinities for Ni(II) without the need for additional chelators (nickelophores) like histidine, suggesting these NiBPs have a unique nickel (‘Ni(II)’) binding mode. However, little is known about the molecular mechanisms underlying the affinity of *Cc*NikZ-II and *Cj*NikZ to nickel and their selectivity for nickel relative to other metal ions. Here, we determined the crystal structure of apo *Cc*NikZ-II by x-ray crystallography and analyzed structures of nine *Cc*NikZ-II homologous proteins by using AlphaFold2 (38, 39). By combining these structural insights with alanine scanning mutagenesis, phylogenetic analyses, and binding site engineering, we characterized the role of the *Cc*NikZ-II residues in Ni(II) binding affinity and engineered higher affinity variants. These results provide new insights into the metal binding determinants of NiBPs, which will enable efforts to further improve their binding affinity and specificity for different metals.

## Results and discussion

### Crystal structure and alanine scanning mutagenesis of CcNikZ-II

*Cc*NikZ-II was recombinantly expressed in *E. coli* with the endogenous signal peptide (30 aa) replaced by a His6-tag and affinity-purified to over 95% homogeneity (Materials and Methods). Purified *Cc*NikZ-II was crystallized using the hanging drop method, and the crystal structure of *Cc*NikZ-II was determined at 2.38 Å resolution by Molecular Replacement (Table S1, PDB 8EFZ). Like other SBPs of Family 5, the *Cc*NikZ-II protomer has a tear-like shape with two α/β domains: domain I with two sub-domains (Ia: residues 1-40, 154-243, 463-526; Ib: 41-153) and domain II (244-462), which encloses a metal binding site located between these lobes (Figure 1A). The *Cc*NikZ-II domain II is connected to sub-domain Ia through a shared anti-parallel pleated β-sheet acting as a rigid hinge based on the apo and holo structure of *Cj*NikZ (PDB 4OET, 4OEV). A Dali search (40) for structurally homologous proteins in the PDB identified over one hundred homologous α/β fold proteins with low overall sequence similarity to *Cc*NikZ-II (17-40% sequence identity). The top structural homologues of *Cc*NikZ-II include *Cj*NikZ from *Campylobacter jejuni* (PDB 4OET, Z-score 51.2, rmsd 1.4 Å, 40% identity), the oligopeptide binding protein AppA from *Bacillus subtilis* (PDB 1XOC, Z-score 42.2, rmsd 2.6 Å, 34% identity), and *Yp*YntA from *Yersinia pestis* (PDB 4OFL, Z-score 40.6, rmsd 2.5 Å, 25% identity). The electrostatic surface charge of *Cc*NikZ-II revealed the central cavity was negatively charged, which strongly suggested the presence of a binding site for positively charged metal ions like Ni(II). Immediately outside the cavity, a ring of neutral surface charge defined a perimeter separating the predominantly positively charged exterior from the cavity (Figure 1B), which we suspect could be a mechanism for limiting adventitious surface binding of Ni(II) via repulsive forces, thus guiding the ions into the cavity.

**Figure 1.**
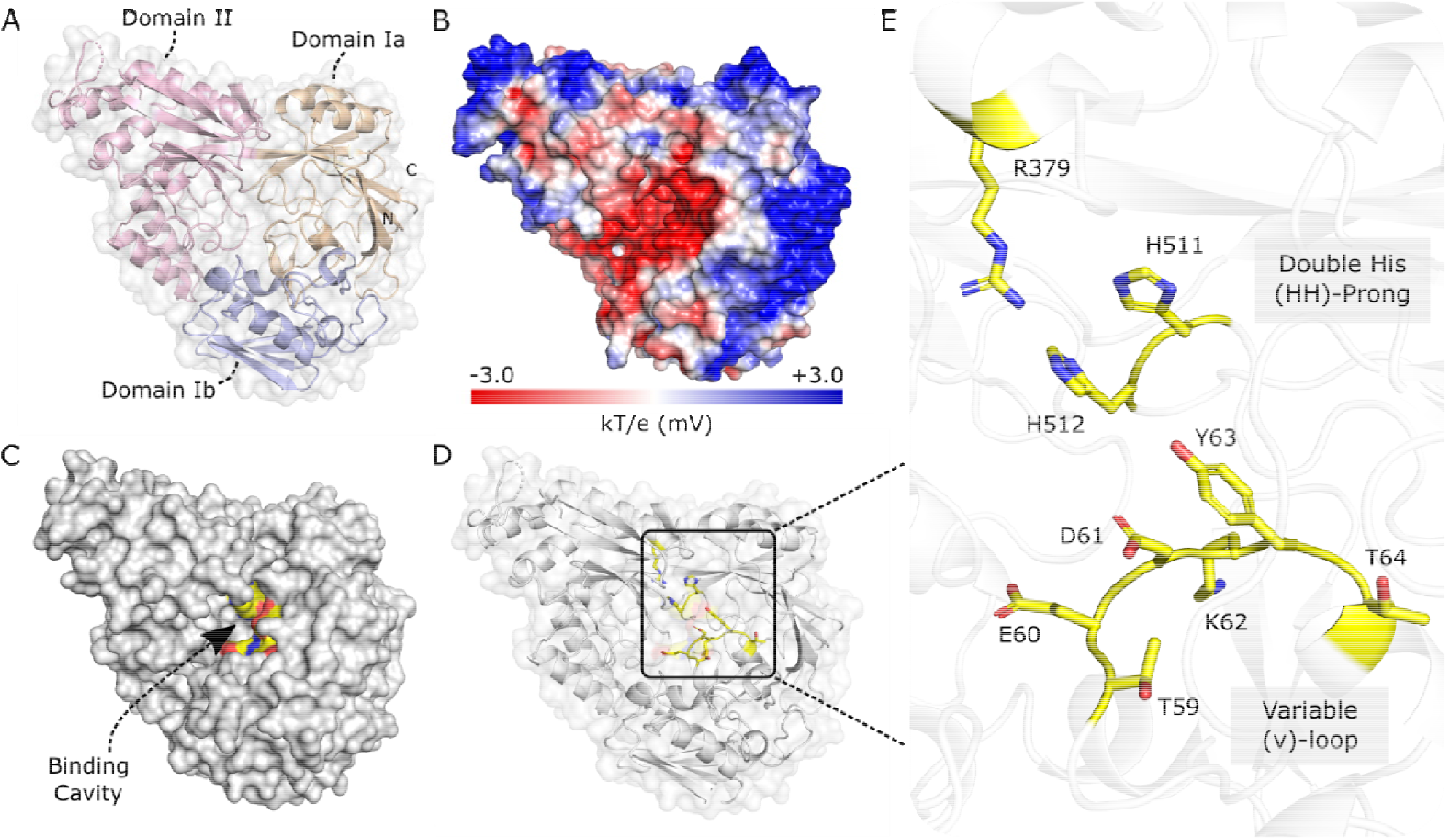
Crystal structure of *Cc*NikZ-II. (A), Overall fold of the *Cc*NikZ-II protomer. The protein is comprised of two domains: the classic bi-lobal domain Ia (tan) and Ib (lavender), and domain II (rose). (B), An electrostatic surface representation generated using an adaptive Poisson-Boltzmann solver. Low kT/e values (red) demark negatively charged areas, and high kT/e values (blue) demark positively charged areas. (C), A surface representation of *Cc*NikZ-II. The binding residues (yellow) are buried deep inside in the core of the protein. (D), Location of the Ni(II) binding residues (yellow) with a (E) close-up view of the *Cc*NikZ-II binding site. The Ni(II) binding site is positioned between the three protein lobes with each lobe contributing residues towards metal binding.

Our recent study of purified *Cc*NikZ and *Cj*NikZ demonstrated these proteins could bind Ni(II) with micromolar affinities without the requirement for adding additional nickelophores such as histidine (41). The structures of *Cj*NikZ complexed with Ni(II) revealed three proteinaceous His residues as direct Ni(II) ligands (His26, His480, and His481), whereas the remaining three ligands to complete the octahedral geometry were provided by a bound free His (PDB 4OEU) or bound oxalate (and a water molecule, PDB 4OEV) acting as additional Ni(II) chelators (21). The *Cj*NikZ Ni(II) binding site also included Arg344 coordinating the bound Ni-chelator (free His or oxalate), as well as several aromatic residues positioned near the bound Ni(II) (Tyr374, Phe380, Trp384, and Phe413). These insights from *Cj*NikZ informed our analysis of the apo *Cc*NikZ-II structure (Figure 1D,E), which revealed the presence of several conserved residues buried deep within the cavity of the protein (Figure 1C), so it was clearly a metal-binding site. The presence of a chloride ion coordinated by the guanidino group of the conserved Arg379 was observed, suggesting that like Arg344 in *Cj*NikZ, this residue can form a salt bridge with the carboxylate groups of a potential small chelator (*e.g*., oxalate or histidine). Similar to His480 and His481 in *Cj*NikZ, we also observed His511 and His 512 positioned such that their imidazole groups pointed into *Cc*NikZ-II’s central cavity. We called these double His motifs in *Cj*NikZ and *Cc*NikZ-II the ‘HH-prongs’. We expected the third His residue homologous to the *Cj*NikZ His26 to be located on a loop near the entrance of the *Cc*NikZ-II central cavity, but found that it was missing. Superimposition of apo *Cc*NikZ-II and apo *Cj*NikZ (PDB 4OET) revealed the structural conservation of these binding residues’ positions. However, *Cj*NikZ’s flexible loop contained one additional residue (A28) that slightly distorted half the loops’ alignment with one another (Figure 2A,B). Taken together, *Cc*NikZ-II’s central cavity was clearly a Ni(II) binding site as our prior work evidenced (41), but the differences in these loops’ composition (herein called variable loops, or ‘v-loops’) suggested there may be differences in how the two proteins bind to Ni(II).

**Figure 2.**
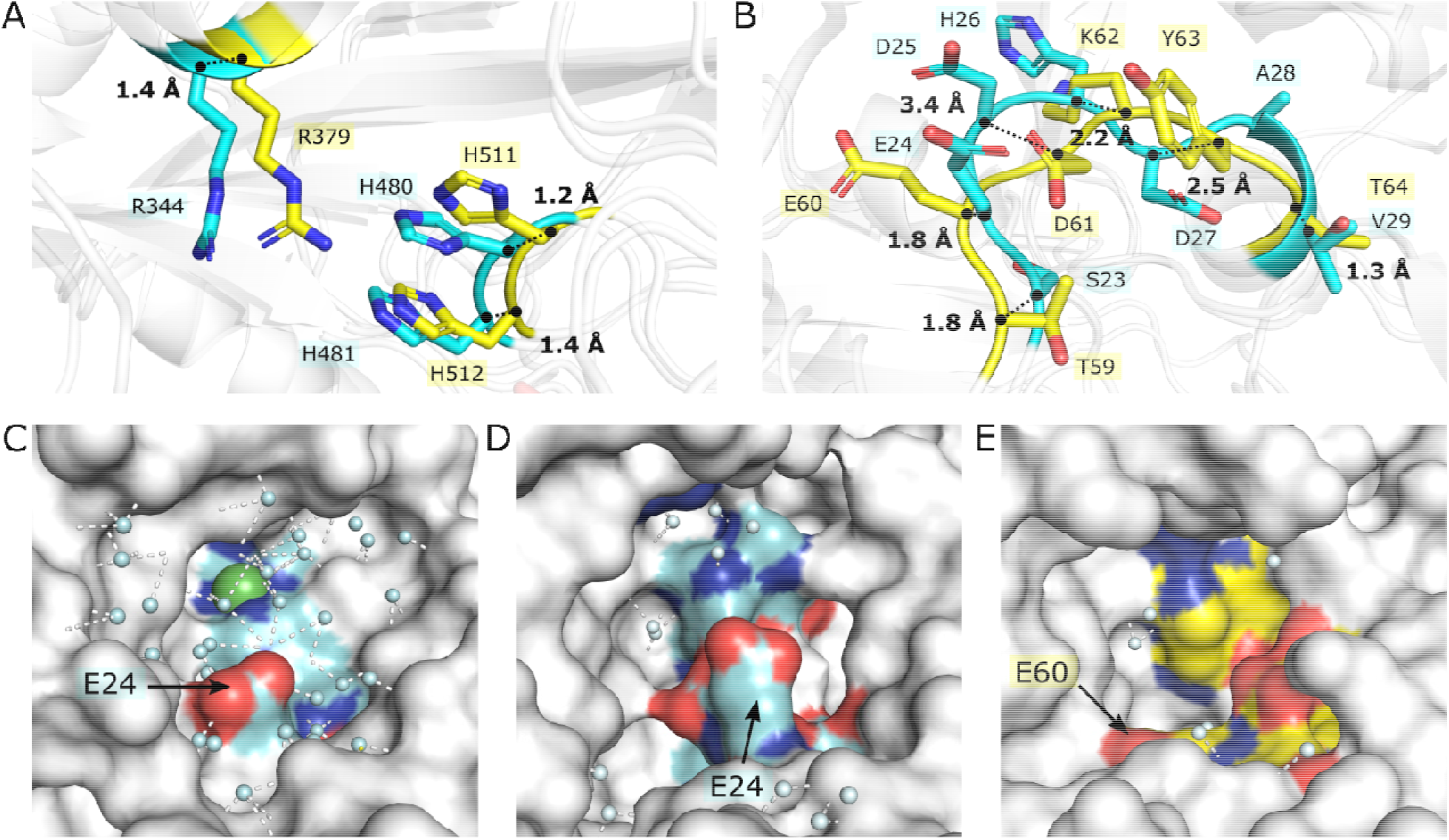
*Structural comparison of* Cc*NikZ-II and* Cj*NikZ*. (A,B), Close-up view of Ni(II) binding site: structural superimposition of apo *Cc*NikZ-II (yellow) and apo *Cj*NikZ (PDB 4OET, cyan) is shown for the upper half (A) and the bottom half (B) of the metal-binding site. Black dotted lines represent the distance between the α carbons for each aligned residue. (C,D,E), Close-up view of the surface of apo *Cc*NikZ-II (C), apo *Cj*NikZ (D), and Ni(II)-bound holo *Cj*NikZ (PDB 4OEV) (E). Water molecules (blue spheres) and their potential polar contacts (white dashed lines) are displayed to show the absence and presence of water networks formed within the binding site. Nickel (green sphere) is shown buried in the site behind the water network.

The v-loop appears to play a dynamic role in Ni(II) binding based on the apo and holo *Cj*NikZ structures. In *Cj*NikZ, the v-loop undergoes a large (>5 Å) movement to reposition its residues (specifically His26) closer to the HH-prong for direct Ni(II) binding. The absence of histidines in *Cc*NikZ-II’s v-loop and thus its unclear role in Ni(II) binding prompted us to use alanine scanning by site-directed mutagenesis to study each binding residue’s impact on *Cc*NikZ-II’s Ni(II) binding affinity (Figure 3, Table S2, Figure S1). We initially targeted residues in the first coordination sphere: Arg379, His511, His512, as well as a double Ala substitution where the HH-prong was substituted (H511A, H512A). The Lys62 of TEDKYT was structurally aligned with the CjNikZ-II v-loop His26, so we hypothesized (‘Hyp1’ for hypothesis 1) it had similar functionality and included it in the initial mutagenesis (K62A). Ala substitution of His511 and His512 each lead to a 2.3-fold and 1.2-fold decrease in binding affinity, respectively, implying that His511 plays a larger role in Ni(II) binding than His512. Surprisingly, the mutant protein K62A had a 4.3-fold improvement in Ni(II) binding affinity. This was likely due to the positive charge of the Lys33 guanidino group at the assay pH (7.2) that would cause it to repel Ni(II), and its bulkiness that may render the v-loop less mobile. The R379A and double H511A/H512A mutant proteins were found to be insoluble when expressed in *E. coli* and could not be purified. Thus, these Ala substitution experiments confirmed the important role of the conserved residues His511, His512, and Lys62 in Ni(II) binding by CcNikZ-II.

**Figure 3.**
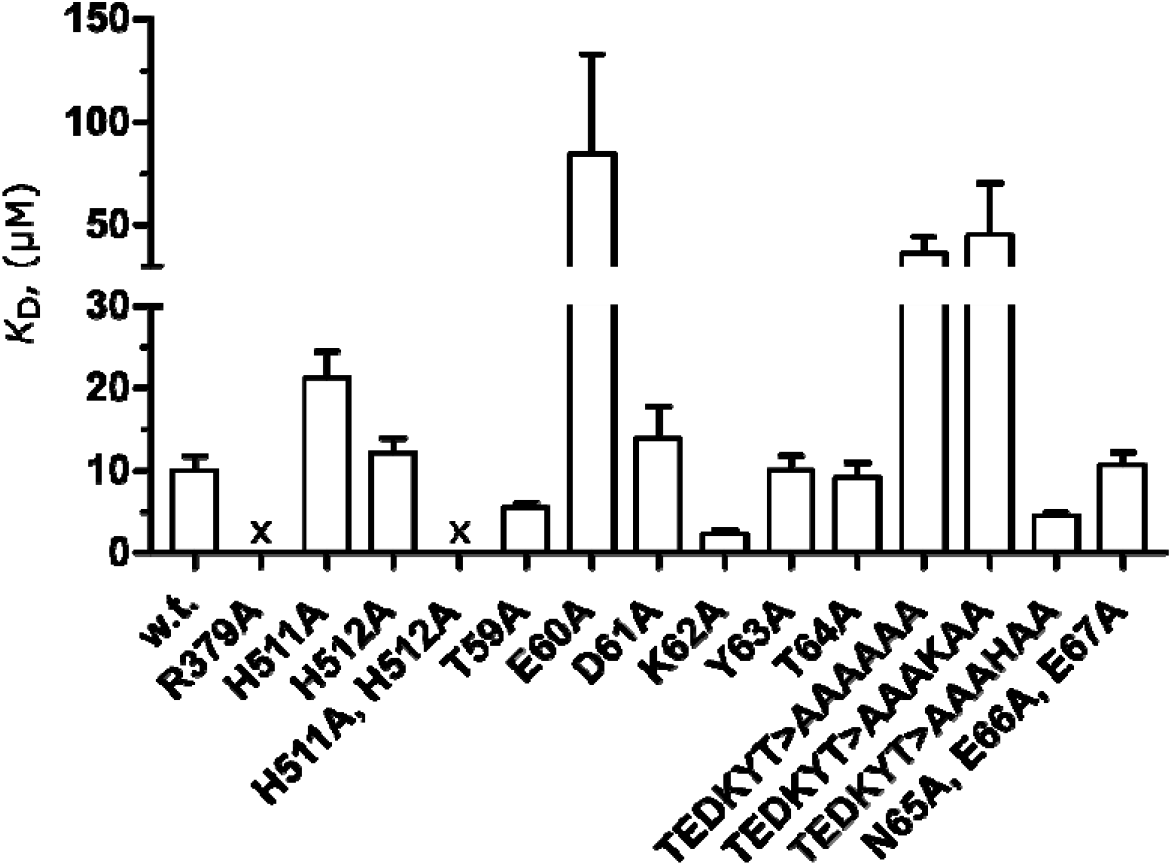
Ni(II) binding affinity of the wild type and mutant *Cc*NikZ-II proteins. Ni(II) binding affinity (*K*_D_) of purified wild type (w.t.) *Cc*NikZ-II and various mutant proteins (single, double, and triple Ala-substitutions and v-loop substitutions). ‘x’ denotes mutant proteins that were generated but could not be purified (insoluble). Experimental triplicates were performed (n=3). Additional numerical data is provided in Table S2.

The unexpected results with the K62A mutant protein led us to probe the role of other residues within the v-loop. We first analyzed the potential polar contacts (*i.e*., hydrogen-bonding) that could be made by water molecules of the original holo *Cj*NikZ structure bound to nickel (PDB 4OEV) and found E24 of its v-loop protruded directly into the water network formed in the binding site (Figure 2C). Compared to apo *Cj*NikZ (PDB 4OET) and our apo CcNikZ-II structures, a water network was not observed in the Ni(II) binding cavities (Figure 2D,E). We therefore hypothesized (Hyp.2) that the Glu60 of *Cc*NikZ-II’s v-loop sequence TEDKYT also played a critical role in its Ni(II) binding affinity by participating in the hydrogen-bonding network. In a subsequent set of mutations (Figure 3), we substituted each residue of TEDKYT with alanine (which included K33A from earlier). E60A confirmed our hypothesis by showing a large 8.4-fold decrease in Ni(II) binding affinity. This was further substantiated with the full substitution of the v-loop by alanines (TEDKYT>AAAAAA) that saw a 3.6-fold decrease in affinity. We did not have sufficient evidence against the hypothesis that K62 was critical to Ni(II) binding affinity at this point, so we believed substituting out all residues in TEDKYT to alanine, except K33 (TEDKYT>AAAKAA), would rescue the binding. On the contrary, we instead saw this mutant protein behaved like TEDKYT>AAAAAA, which agreed with the prior K33A observation. Remarkably, replacing the lysine in this mutant protein with a histidine (TEDKYT>AAAHAA) improved the binding affinity 2.2-fold over the wild-type TEDKYT. The end of *Cc*NikZ-II’s v-loop is connected to a small α-helix with the sequence NEE. Given its spatial distance from H511 and H512, we expected no change in binding affinity upon alanine substitution, which we confirmed with the triple mutant protein (N65A, E66A, E67A) that showed little change in affinity. Together, this pointed to the necessity of E60 and the advantage of having a histidine closely adjacent to it, leading to our next hypothesis (Hyp.3) that core v-loops containing both a glutamic acid and histidine within close proximity near the front of the loop would confer the highest Ni(II) binding affinities amongst homologues. Milder changes in affinity observed in the D61A (1.3-fold decrease in affinity) and T59A mutant protein (1.8-fold increase in affinity) were alone insufficient for elucidating their role. We also noted that the Y63A and T64A mutant proteins showed little change in affinity, suggesting residues in these positions may contribute less to Ni(II) binding affinity. To further understand how v-loops can impact Ni(II) binding affinity, we next used a phylogenetic approach to analyze the diversity of v-loops found in *Cc*NikZ-II and *Cj*NikZ’s NiBP family.

### Phylogenetic analysis of NiBPs and v-loop sequence anatomy

In prior work, Lebrette *et al*. (2014), created a global NiBP phylogeny by performing a BlastP search on NCBI using eight experimentally studied NiBPs (*Ec*NikA, *Yp*YntA, *Vp*NikA, *Hh*NikA, *Cj*NikZ, *Sa*NikA, *Sa*CntA, *Bs*NikA) as seeds, then manually extracting the first 1,000 homologs displaying the highest similarity for each seed, followed by a decrease in the redundancy of the 8000 sequences through several trimming routines to obtain a representative library with 3276 sequences (21). They observed high sequence similarity between NiBPs and PepBPs, so they proposed that the substrate binding promiscuity between the NiBPs and PepBPs resulted from a shared evolutionary history. We created a global phylogeny comprised of members from both groups using InterPro where the SBPs are organized according to the Tam-Saier Classification (Figure 4). We first decreased the redundancy of the InterPro family IPR030678 using the CD-HIT algorithm to obtain a representative library of 4561 NiBP and PepBP sequences (42, 43). We then used FastTree to generate a phylogenetic tree from a MAFFT alignment to obtain a representative library (Figure 1) (44). The difference in these two approaches is Lebrette *et al*.’s approach introduced bias due to selection of the pool of sequences of experimentally characterized NiBPs for the initial BlastP search. This means that alternative modes of binding in extant NiBP and PepBPs which could have promiscuity for Ni(II) binding may have been under-represented and overlooked. By using InterPro, we captured a fuller diversity of NiBP/PepBPs and thus renewed some observations made by Lebrette *et al*. (2014) while also observing major differences (Supplementary Data File 1, 2).

**Figure 4.**
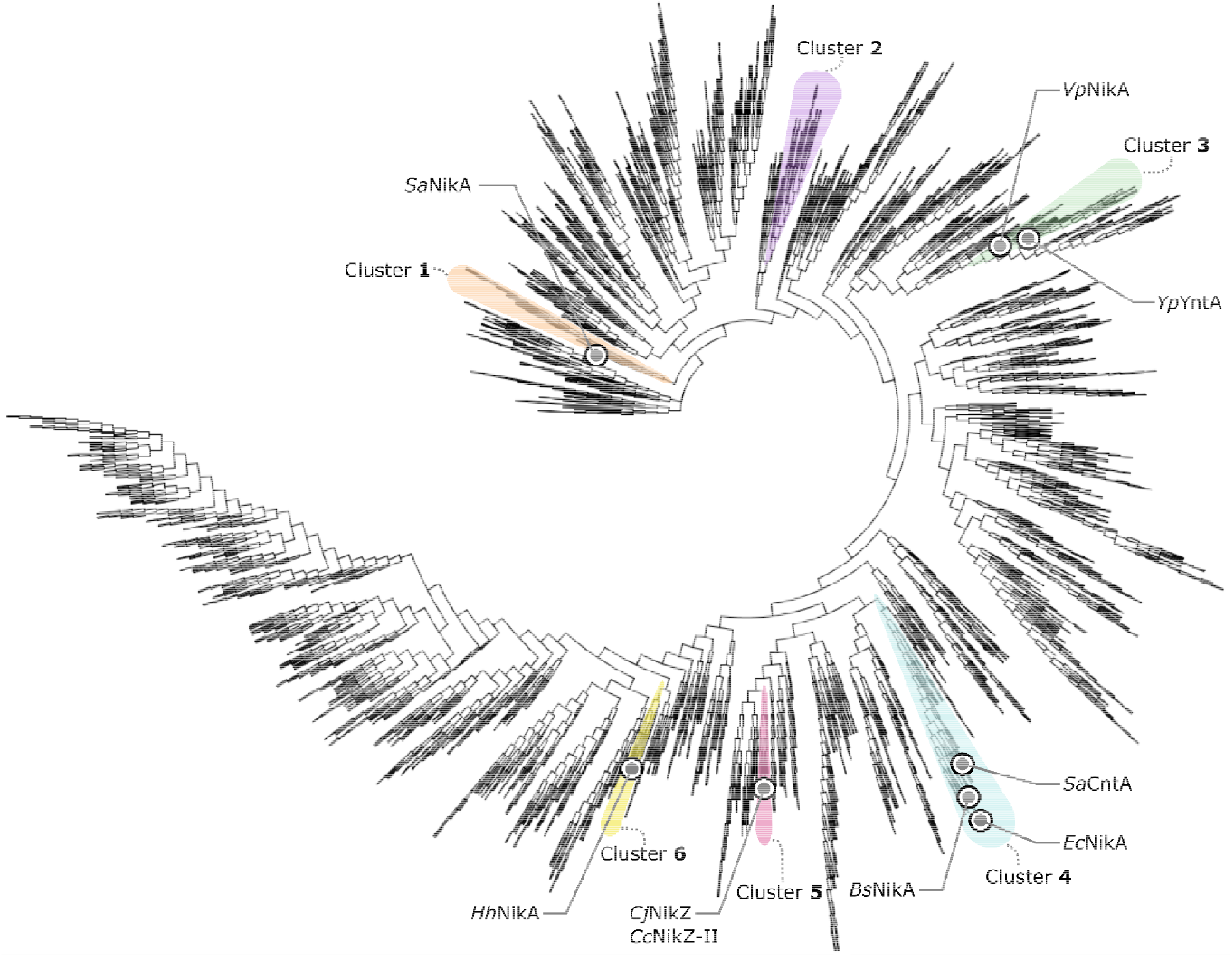
Phylogenetic analysis of NiBPs. The representative library was created by applying the CD-HIT algorithm to the InterPro family IPR030678 such that sequences with 60% redundancy were removed. From this, the phylogenetic tree representation of the representativ library built using FastTree on Geneious is shown. Experimentally characterized NiBPs (grey circles) are mapped onto the tree with coloured spheres demarking their phylogenetic branch.

Similar to Lebrette *et al*. (2014), we found that *Vp*NikA and *Yp*YntA cluster together (Figure 4, Green), while *Sa*CntA, *Ec*NikA, and *Bs*NikA cluster together (Figure 4, Blue). In contrast to Lebrette *et al*. (2014), *Sa*NikA (Figure 4, Cluster 1) clusters further away from *Vp*NikA and *Yp*YntA (Cluster 3), and closer to the root. *Sa*NikA follows the third binding strategy outlined by Lebrette *et al*. (2014) where no direct contact by the protein is made with the nickel ion. Instead, proteinaceous arginine and aromatic residues are used to interact with the free histidine amino acids that are complexed with nickel. *Sa*NikA also lacks the proteinaceous histidines found in other NiBPs that follow the outlined first and second binding strategies where some degree of direct contact (*viz*. coordinate covalent bond) is made with the nickel ion. This, combined with the positioning of *Sa*NikA in our phylogenetic tree, suggests that ancestral proteins of the IPR030678 family may have been PepBPs originally, but because peptides are strong complexing agents for nickel, some members of the family may have evolved specificity for nickel and nickel complexes. More strikingly different, *Cj*NikZ (Cluster 5) appeared closer to *Ec*NikA homologs (Cluster 4) than *Vp*NikA and *Yp*YntA (Cluster 3) in Lebrette *et al*. (2014), suggesting *Cj*NikZ shared a common ancestor with *Ec*NikA despite having only 25% identity with each other. This difference in positioning of *Cj*NikZ may be attributed to the addition of PepBPs to build the global phylogeny wherein *Vp*NikA and *Yp*YntA’s overall structures have higher similarity to PepBPs with ancestry closer to the root.

We focused our attention on the ‘close homologues’ of Cluster 5 (Figure 4) where both *Cc*NikZ-II and *Cj*NikZ can be found. By searching for the HH-prong motif ‘GHHG’ across the representative library, we also found proteins in Cluster 2 that appeared to be ‘distant homologues’ of *Cc*NikZ-II. Upon closer inspection, 21/24 members (88%) of Cluster 5 contained the GHHG motif, whereas only 8/78 members (10%) of Cluster 2 contained the HH-prong in a relatively similar position, but not the exact GHHG motif (*viz*. the glycines were replaced with other small hydrophobic residues). We extracted sequences for nine close homologues and four distant homologues based on their uniqueness within their local phylogenies, and whether we already cloned them for expression. After combining these sequences with *Cc*NikZ-II and *Cj*NikZ, their sequence alignment revealed three observations (Figure 5A): 1) the positions of the arginine and HH-prong were highly conserved, and 2) some of the distant homologues had distinctly different sequences in *Cc*NikZ-II’s v-loop region, whereas 3) the distant homologue from *Nocardiopsis gilva* and the close homologues’ sequences in this region did share similarities, but the exact positional conservation was unclear.

**Figure 5.**
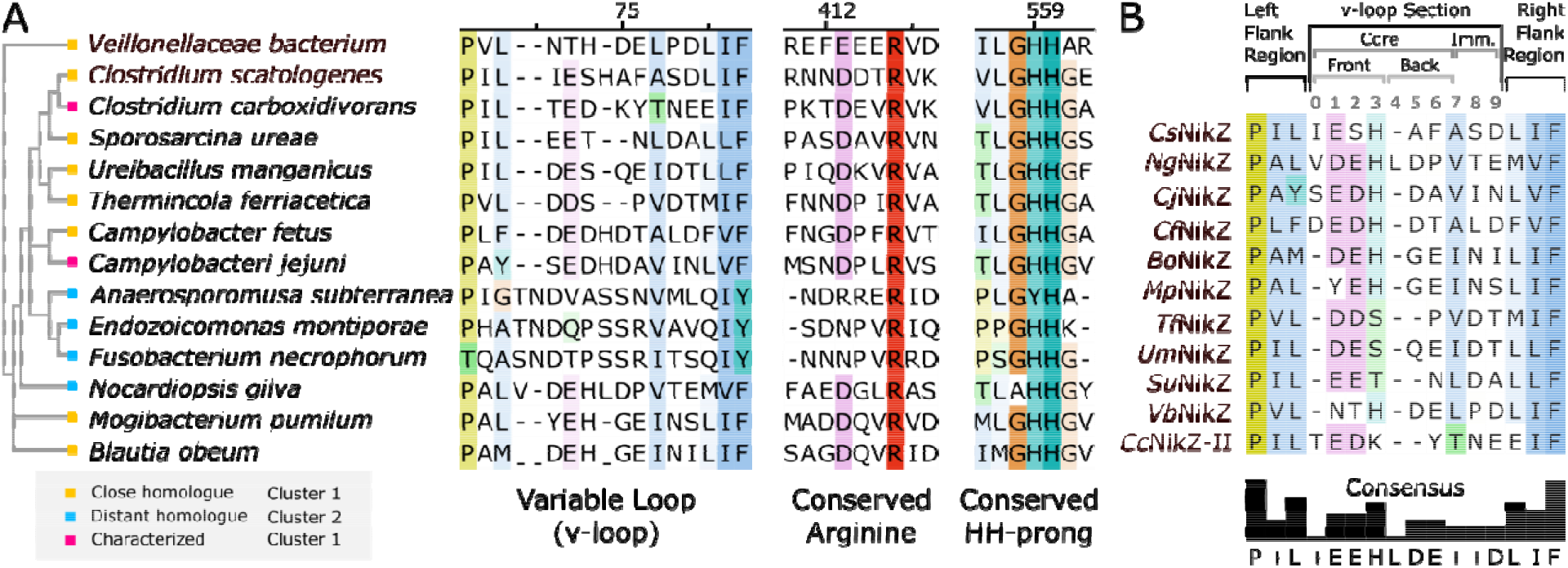
The sequence motifs of selected NiBPs involved in Ni(II) binding. (A) Ordered according to their local phylogeny, several sequences of close (yellow) and distant (blue) homologues of already characterized *Cc*NikZ-II and *Cj*NikZ (pink) are aligned using MAFFT with coloring based on ClustalX at 25% conversation threshold. Three regions are shown: the variable loop (v-loop), the conserved arginine, and the conserved double histidine (HH-)prong with the numbers on the top indicating (or add the CcNikZ-II residue numbers). (B) Ordered according to their local phylogeny, sequences of selected members of the wild-type NiBP set are aligned and coloured as before. Naming is based on their microorganism origins (*cf*. Table S3). The v-loop anatomy is shown (top). The v-loop section is between the left flank region and right flank region, which are strongly conserved. The v-loop section is comprised of the flexible core subsection (core v-loop) and the immobilized subsection (an α-helix). The core v-loop possesses a front and back to demark their different functions. The consensus sequence (bottom) is displayed, where ‘+’ indicates the positions that do not have clear conservation.

We therefore kept the sequence for *N. gilva* and removed the other distant homologues to create this study’s wild-type NiBP set. We gave new names for the sequences based on their source microorganism and their anticipated ability to bind to nickel like *Cc*<u>NikZ</u>-II and *Cj*<u>NikZ</u> (*e.g*., *Cs*NikZ, from *<u>C</u>lostridium <u>s</u>catologenes*, Table S3). A re-alignment of the sequences revealed critical details about the v-loop anatomy (Figure 5B, Figure S2). The v-loop section was flanked by two strongly conserved consensus sequences: P+L for the left flank region and LIF on the right flank region (‘+’ here indicating valine, isoleucine, and alanine). The v-loop section itself was 5-8 residues long and comprised of the flexible core and immobilized sub-sections. The flexible core (herein ‘core v-loop’) contains a front and back to delineate their different roles with respect to Ni(II) binding affinity. The last residue of the core v-loop and the first residue of the immobilized subsection are the same (position 7) to reflect the unclear boundary between the two subsections. Apart from *Cs*NikZ, *Ng*NikZ, *Cj*NikZ, and *Cf*NikZ that have an addition residue at position 0, the sequence EEH (positions 1-3) is the consensus sequence for the front of core v-loop. The majority of the homologues had either aspartate or glutamate at positions 1 and 2, followed by a histidine at position 3. This observed conservation across the wild-type NiBP set pointed to our prior hypothesis (Hyp.3) that the glutamate followed by a nearby histidine in the front of the core v-loop conferred higher Ni(II) binding affinities. Before determining *K*_D_ values for the wild-type NiBP set to test Hyp.3, we created the engineered *Cc*NikZ-II variant set to simultaneously test another hypothesis (Hyp.4) that core v-loops had a degree of modularity whereby loop exchanges from high-affinity homologues could improve *Cc*NikZ-II’s Ni(II) binding affinity.

### Uncovering core v-loop modularity and unique homologue properties

The core v-loops shown in Figure 5B differed in length (5-8 residues) and composition. Rather than simply replace the 6-residue TEDKYT core v-loop of *Cc*NikZ-II with the homologues’ core v-loops, which could confound the effects of their length and composition on Ni(II) binding affinity, we first assessed the placement of the homologue’s core v-loops relative to their binding sites. This was to understand how the core v-loop lengths affected the physical placement of the residues in its composition. Since AlphaFold2 generated a more accurate prediction of apo *Cc*NikZ-II’s structure compared to SWISS-MODEL (Figure S3), we used AlphaFold2 for modelling 3D structures of selected *Cc*NikZ-II homologues, which are shown in Figure 6 with highlighted the 6-residue sequences that were selected for loop exchange with the *Cc*NikZ-II core v-loop (TEDKYT). These selections were primarily based on considerations combining Hyp.3 (*viz*. the critical positioning of a glutamate at the front of the core v-loop) and structural confirmation that the residues’ positions were similar to *Cc*NikZ-II and *Cj*NikZ’s core v-loops (Figure 1D). For example, *Bo*NikZ possessed a glutamate (E22) with a histidine (H23) immediately adjacent, so the remaining four residues were chosen to situate E22 at the front of the core v-loop while preserving its second-from-the-start position: DEHGEI. The structure shown in Figure 6A helped confirm this selection since the final isoleucine (I26) was positioned near the start of the immobilized subsection, similar to *Cc*NikZ-II’s T35 (Figure 1D). The *Cc*NikZ-II engineered variant based on this selection was thus named *Bo*>*Cc* to denote the presence of DEHGEI in lieu of (>) TEDKYT. For *Vb*NikZ and *Tf*NikZ, a glutamate was not present in the front of the core v-loop, so we simply used the first six residues following their left flank regions.

**Figure 6.**
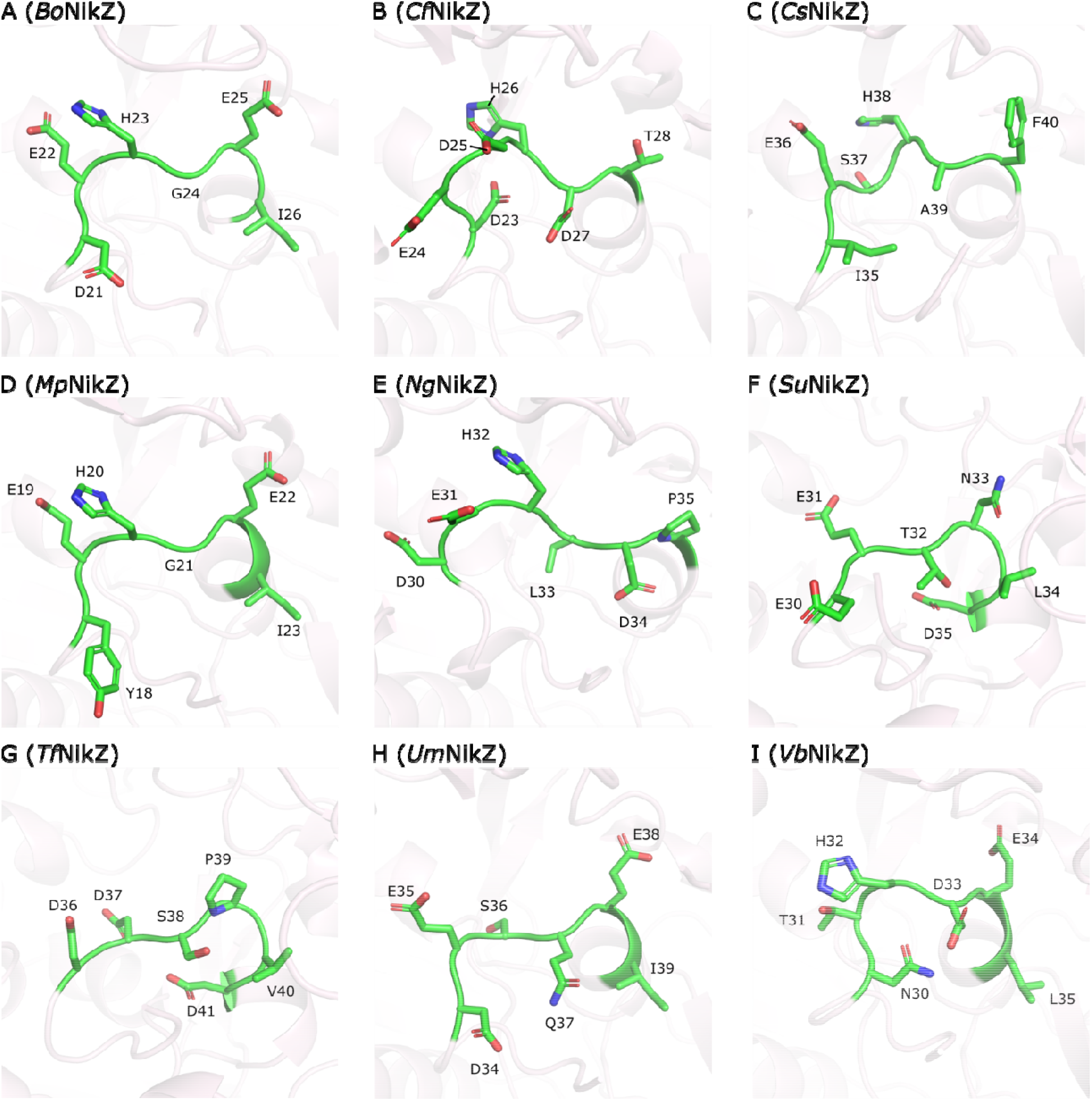
Structural models of the *Cc*NikZ-II homologues’ core v-loops by AlphaFold2. Each panel displays the segment of the core v-loop (green) that is used to substitute *Cc*NikZ-II’ core v-loop sequence TEDKTY. Residues are numbered according to their respective sequences. (A), *Bo*NikZ (DEHGEI). (B), *Cf*NikZ (DEDHDT). (C), *Cs*NikZ (IESHAF). (D), *Mp*NikZ (YEHGEI). (E), *Ng*NikZ (DEHLDP). (F), *Su*NikZ (EETNLD). (G), *Tf*NikZ (DDSPVD). (H), *Um*NikZ (DESQEI). (I), *Vb*NikZ (NTHDEL).

The Ni(II) binding affinities were next determined for the wild-type NiBP and engineered variant set (Figure 7, Table S2, Figure S4-S6). Among the homologues, *Cc*NikZ-II had the lowest Ni(II) binding affinity (Figure 7), which we explained according to Figure 7B where *Cc*NikZ-II’s v-loop section did not have a histidine following the front’s glutamates and aspartates in position 1 and 2, and was the only member to have a hydrophilic residue (T64) at position 7. In the same breath, this meant all other homologues had higher Ni(II) binding affinities. Ranking the homologues according to affinities (Figure 7B) provided evidence supporting Hyp.3 where the highest Ni(II) binding affinities corresponded to homologues with glutamates at position 2 and histidines at position 3. The back of the core v-loops (positions 4-9) did not exhibit notable trends apart from position 7, further supporting the earlier notion that they may play other roles that do not impact Ni(II) binding affinity. Among the engineered variants (Figure 7C), *Cc*NikZ-II again had the weakest Ni(II) binding affinity, which generally supported Hyp.4 in that all wild-type NiBPs had higher affinities, and their cognate engineered varaints also had higher affinities than wild-type *Cc*NikZ-II. Ranking the engineered variants (Figure 7D) further substantiated Hyp.4 where the top five proteins for both the sets were based on the same core v-loops from *Bo*NikZ, *Cf*NikZ, *Cj*NikZ, *Mp*NikZ, and *Ng*NikZ. However, the orders of the rankings were different, which may reflect contributions from the v-loop length or other residues found in the homologues’ binding sites that differ from *Cc*NikZ-II’s binding site (*vide infra*). We also noted that aspartates and glutamates at position 4 were also a common feature of the top five engineered variants (Figure 7D). Based on the physical placement of *Cc*NikZ-II’s Y63 where residues at position 4 would exist (Figure 1D), it is plausible that aspartates and glutamates are better at contributing to a water network than tyrosine, thus improving Ni(II) binding affinity (Figure 1E).

**Figure 7.**
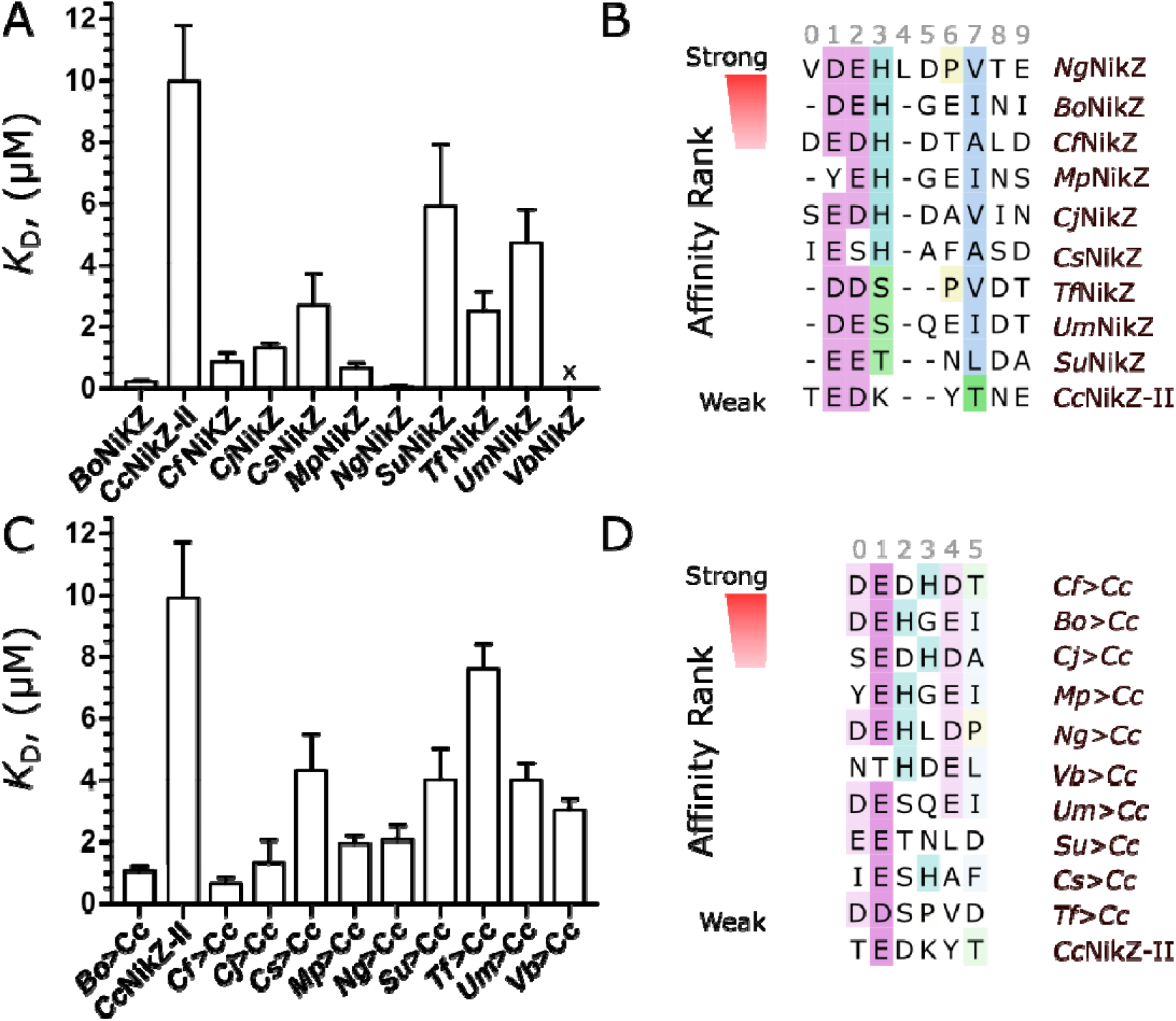
Ni(II) binding affinities of purified wild type and mutant NiBPs. Ni(II) binding affinities and ranking of purified wild type NiBPs (A, B) and engineered CcNikZ-II variants with different v-loops (C, D). The position numbering (grey) is the same as the numbering in Figure 5B). Sequences are aligned based on MAFFT and coloured according to ClustalX with a 25% conservation threshold. The position numbering is not the same as the numbering in Figure 5B. ‘x’ denotes when a binding curve could not be generated (no observable change in fluorescence). Experimental triplicates were performed (n=3). Additional numerical data is provided in Table S2.

While K62A and *Cf*>*Cc* exhibited the largest improvements in *Cc*NikZ-II’s Ni(II) binding affinity, we also discovered two homologues with interesting properties. First, *Ng*NikZ was determined to have a sub-micromolar *K*_D_ value that should be carefully interpreted as an upper limit to its true *K*_D_ value since it was less than the concentration of the protein in the ITFQ assay (45). This exceptionally high-affinity binding may be due to its origins in *Nocardiopsis gilva* YIM 90087, a gram-positive species discovered in hypersaline environments in China (46). Li *et al*. (2006), determined *N. gilva* could grow in 18% (w/v) NaCl, with an optimum range around 5-8%. In previous work, we showed how an increasing the concentration of NaCl (up to 1 M) decreased *Cc*NikZ-II’s Ni(II) binding affinity, which suggests NGI may have evolved to have exceptionally higher Ni(II) binding affinity to overcome the presence of salts in order to capture nickel for the cell (37). The importance of reaching higher affinities in metal recovery application is explored in our other work (unpublished) where our model showed how increasing the affinity of a NiBP for a target metal could allow it to sequester more of that target metal from a complex multi-metal solution. Additional characterization of *Ng*NikZ is thus warranted. Second, we measured melting temperatures (*T*_m_) for all members of the Ala substation proteins and proteins from the two sets to assess the quality of their folding prior to the ITFQ assay (Figure S7). We found that *Tf*NikZ had the highest *T*_m_ at 76 °C, which likely reflected its origins in hydrothermal springs of Japan, where its source microorganism *Thermincola ferriacetica* was found to grow between 45-70 °C and most optimally at 60 °C (47). *Tf*NikZ’s heat resistance property may be useful for higher temperature applications. Overall, these mutations and traits highlight the importance of developing higher-throughput methodologies to screen for and study metalloproteins with exceptional properties. Since we found the rankings of the wild-type NiBP and engineered variant proteins with the highest Ni(II) binding affinities were not the exact same, potentially due to differences in the proteins beyond the v-loop, it prompted us to explore the use of automation to test the hypothesis (Hyp.5) that core v-loops from wild-type NiBPs could transfer their native metal specificity preferences (herein ‘promiscuities’) to *Cc*NikZ-II. Here, metal promiscuity refers to the set of metals that a protein can bind to. The more specific the protein the smaller the set of metals it can bind to, which is important for target metal recovery from multi-metal solutions.

### Metal promiscuity screening by automated assays

To test Hyp.5, members of the two sets needed to be systematically titrated with different concentrations of metals, and any fluorescence changes would need to be computed to determine whether metal promiscuities were similar between a wild-type NiBP and its cognate engineered variant (*e.g*., *Cf*NikZ and *Cf*>*Cc*). If their metal promiscuities were similar, we expected to see matching ‘hit’ profiles (*e.g*., the set of metals with significant binding to *Cf*NikZ were the same set for *Cf*>*Cc*). The large number of replicates required (>1000 wells) prompted us to develop an automated screening workflow using the Tecan Freedom EVO liquid handler coupled with the same Tecan microplate reader used in prior experiments. The workflow is divided into four stages (Figure 8A): Prepare, Test, Analyze, and Visualize. In the Prepare stage, the liquid handler and microplate reader are programmed to move and perform experiments with 96-well opaque microplates. In the Test stage, the proteins are first dispensed into plates and equilibrated to measure the blank fluorescence (*F*_0_). Then, the liquid handler dispenses activity buffer (10 mM HEPES, pH 7.2) as the negative control (no metal), and an assortment of metals (Ni^2+^, Co^2+^, Cu^2+^, Zn^2+^, Mn^2+^, Al^3+^, Pb^2+^, Cd^2+^, La^3+^, Eu^3+^, and Yb^3+^) to final concentrations of 100 nM and 100 μM in different wells. Finally, the samples are equilibrated to measure the fluorescence (*F*). To move to the Analyze stage, the *F*_obs_ is calculated (*F-F*_0_) for each metal-protein pair and a Python script then performs a statistical test to determine whether a significant metal binding event (hit) occurred at 100 nM and 100 μM of the metals. Depending on the p-value of the hit, a gradation of red blocks was assigned that were then plotted in the Visualize stage as a heatmap.

**Figure 8.**
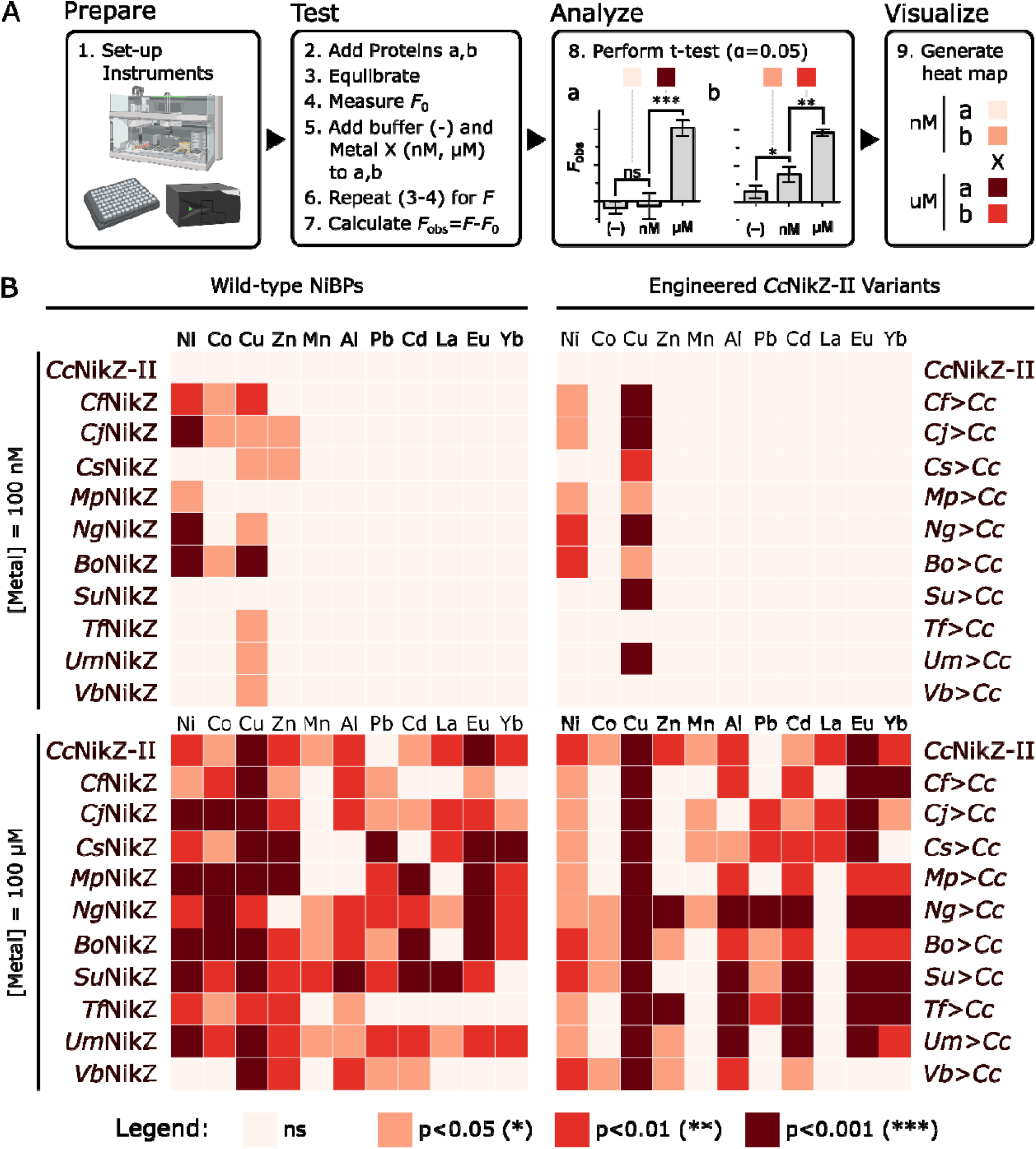
Screening of purified wild type and engineered NiBPs for binding different metal ions. (A) Summary of an automated screening workflow comprised of four stages: Prepare (equipment set-up), Test (ITFQ assay), Analyze (compute statistics with Python), and Visualize (generate heatmap with Python). Proteins (a and b) are titrated with activity buffer as a negative (-) control, and metal solutions (X) to a final concentration of 100 nM and 100 μM. Based on the p-value computed between the average *F*_obs_ of the negative control and 100 nM, and between the 100 nM and 100 μM metal, blocks with gradations of red are applied to each metal-protein pair. If the block is colored red (not light link), it is considered a nM-hit or μM-hit depending on the metal concentration. Darkness of the blocks represent the p-values (legend). (A), Binding of different metal ions by purified NiBPs: heatmap presentation for nM-hits and μM-hits. Members of the wild-type NiBP set (left column) and the engineered variant set (right column) equilibrated with 100 nM (top row) and 100 μM (bottom row) for eleven different metal ions (prepared using chloride salts). Experimental triplicates (n=3) were performed for the negative (-) control and 100 nM metal solutions, and duplicates for the 100 μM solutions (due to limited space on 96-well plates).

Qualitatively, the darker the block, the more statistically significant the hit due to larger changes in fluorescence between the negative control and 100 nM samples (a nM-hit), and between the 100 nM and 100 μM samples (a μM-hit). The reader should take caution in comparing the darkness of different blocks. It is inadvisable to compare the block’s darkness for a metal-protein pair relative to other blocks for the same protein, but a different metal at any concentration, since metal-protein pairs will often have their own unique *F*_max_ values that are mutually exclusive from their *K*_D_ values (*e.g*., *Cc*NikZ-II with 100 μM nickel and 100 μM europium). Similarly, comparing a block’s darkness for a metal-protein pair relative to other proteins, but for the same metal at any concentration, bears the same caveat. The best use of the block’s darkness is to compare a metal-protein pair to blocks for the same protein with the same metal at a difference concentration, which can be used to assess whether the metal-protein pair exhibited nM or μM affinity (*e.g*., *Cc*NikZ-II with nickel, which only has a μM-hit and therefore does have nM nickel affinity), and whether saturation was reached in the concentrations tested (this was not observed in our screen, but an example would be the presence of an nM-hit for a metal-protein pair that did not have a corresponding μM-hit for the same metal).

We applied this workflow to purified wild type and engineered NiBPs from this study (Figure 8B). There were three observations we expected, serving as positive controls for this metal promiscuity screen. First, we expected copper to have hits across all proteins since it forms the most stable complexes according to the Irving-William series of divalent metal-binding preferences: Mn(II) < Fe(II) < Co (II) < Ni(II) < Cu(II) > Zn(II) (48). Indeed, we found copper had the greatest number of nM- and μM-hits with p < 0.001. Second, we expected μM-hits for nickel across all proteins given the previously discussed observations wherein proteins from the two sets (Figure 7A,B) all had tighter Ni(II) binding than *Cc*NikZ-II, which we observed in this screen except for *Vb*NikZ. However, this agreed with a prior observation (Figure 7) where a binding curve could not be generated for *Vb*NikZ with nickel. Lastly, we expected the proteins with the highest Ni(II) binding affinities (Figure 6B) to have a higher likelihood of manifesting as nM-hits. *Cf*NikZ, *Bo*NikZ, *Cj*NikZ, *Mp*NikZ, and *Ng*NikZ (herein considered ‘tight-binders’) displayed nM-hits for nickel. Their cognate engineered variants also had nM-hits. These confirmations enabled further analysis.

While the tight-binders all had nM-hits, they did not have the same nM-hit profiles. Most notably, *Cj*NikZ could bind to 100 nM Zn(II), which other tight-binders could not. *Cf*NikZ and *Bo*NikZ also shared *CjNikZ*’s ability to bind 100 nM Co(II), whereas *Mp*NikZ and *Ng*NikZ had more specific nM-hit profiles. *Cc*NikZ-II displayed no nM-hits for any metal, so we assessed Hyp.5 for the transferring of nM-hit profiles from the tight-binders to *Cc*NikZ-II (*viz*. their cognate engineered variants) and found nM-hits for nickel and copper could be transferred but could not be transferred for zinc and cobalt. This indicated the residues in *Cc*NikZ-II’s binding site combined with the higher Ni(II) binding affinity core v-loops from tight-binders could, unexpectedly, improve nickel specificity relative to the cognate homologue. For example, *CjNikZ* had nM-hits for Ni(II), Co(II), Cu(II), and Zn(II); whereas *Cj*>*Cc* only had nM-hits for Ni(II) and Cu(II), which exemplifies a narrowing (*viz*. improvement) of the nM-hit profile specificity. With respect to Hyp.5, this provided some evidence to claim v-loops can confer at least partial nM-hit profile specificity. However, from a protein engineering perspective, this novel partiality is important if the specificity becomes narrower in favour of the target metal (*i.e*., nickel). From this perspective, *Cf*>*Cc, Cj*>*Cc*, and *Bo*>*Cc* exhibited higher nickel specificity than their cognate wild-type NiBP and are examples of how core v-loop exchanges can simultaneously improve affinity *and* specificity.

The μM-hit profiles were notably more diverse for the wild-type NiBPs, which we understood as a reflection of their different structures and origins. While we found near-unanimous μM-hits for nickel, cobalt, copper, and zinc (herein ‘high-affinity metals’) across the wild-type NiBPs, the μM-hits for the other metals (herein ‘low-affinity metals’) was more sporadic. Although these homologues possessed conserved features (*e.g*., arginine, HH-prong, the v-loop composition for tight-binders) contributing to their shared Ni(II) binding abilities, we contend (Hyp.6) the differences in their second coordination sphere and overall protein matrix likely conferred different abilities to discern competing low-affinity metals that could exist in the environment. This notion is frequently discussed in the literature (5, 49) and was supported by our prior observations where 1) the order of the Ni(II) binding affinity rank (Figure 7B,D) for the tight-binders and their cognate engineered variants did not completely match, and 2) the nM-hit profiles could only partially be transferred to *Cc*NikZ-II. Here, we found additional evidence for Hyp.6 in the engineered variant set where it was more likely to observe μM-hit profiles for low-affinity metals that were the exact or nearly the same (*Cf*>*Cc, Mp*>*Cc*; *Bo*>*Cc, Su*>*Cc, Tf*>*Cc, Ng*>*Cc, Um*>*Cc*; *Cj*>*Cc, Cs*>*Cc*). In theory, these engineered variants have the same *Cc*NikZ-II structural template, aside from the exchanged 6-residue core v-loop, meaning the determinants for their promiscuity were kept relatively constant compared to members of the wild-type NiBP set.

With respect to Hyp.5, we found some of the tight-binders’ cognate engineered variants (*Ng*>*Cc* and *Bo*>*Cc*) exhibited μM affinity to cobalt and zinc, whereas other tight-binders did not. This introduced additional nuance to Hyp.5 where the specificity conferred through some core v-loops may not hold at higher metal concentrations. Future studies could explore the ‘persistence’ of the specificity as a different protein engineering metric, where higher persistence (narrow specificity despite higher metal competition) would be desirable for more robust metal recovery from multi-metal solutions. Finally, the μM-hits observed for most protein of the sets with the rare earth elements (REEs) lanthanum, europium, and ytterbium are promising. Protein engineering efforts may alter *Cc*NikZ-II’s specificity towards REEs via mutagenesis of His482, His483, and the v-loop to accommodate REEs’ preference for Class A ligands (O-donor groups) with hard properties. Taken together, this metal promiscuity screen raises questions about how one knows that an SBP is a Ni(II) specific SBP, and not a Cu(II) specific SBP (for example) due to the Irving-William series that generally dictates stronger affinities between proteins and Cu(II) *in vitro*.

Cells have evolved intracellular systems to correctly metalate proteins with low-affinity metals like Mn(II) and Fe(II) (4, 50, 51), but how could it be possible for there to be periplasmic Mn(II)-specific SBPs (*e.g*., PsaA from *Streptococcus pneumoniae*) despite their bottom rank in the Irving-William series? It suggests there is likely a mechanism that cells have evolved to ensure active Mn(II) acquisition by Mn(II) specific ABC importers. (52, 53). Heddle *et al*. (2003) have hinted at this by calculating there to be ~120 μM of an NiBP (*e.g*., *Ec*NikA) in the periplasmic space of an *E. coli* cell. With a *K*_D_ ≈ 10 μM affinity for Ni(II), up to a 13-fold reduction in Ni(II) efflux out of the periplasm could be achieved (54). So, although NiBPs like *Cc*NikZ-II and its homologues can bind to other metals with appreciable nM and μM affinity, their protein concentrations relative to the concentration of their target metal and their affinity for it determines their collective ability to capture the target metal. The quantitative relationship describing this interplay *in vivo* has not been studied yet, to our knowledge, but may help uncover why the v-loop is not conserved across members of Cluster 2 and 4. Indeed, this is an area we are actively exploring since these principles could apply to metal separation biotechnologies where these NiBPs are used *in vitro*. Specifically, how an NiBP’s affinity and specificity for Ni(II) is related to its rate of Ni(II) transport (*viz*. capture) is an intriguing question. It is possible that further engineering efforts to improve NiBP affinity and specificity for Ni(II) may yield variants with supreme Ni(II) capturing performance.

## Conclusions

Overall, our approach to understanding the v-loop’s contribution to metal binding combined a bioinformatic analysis with biochemical and structural characterization to address six hypotheses. Based on superimposition of our apo *Cc*NikZ-II’s structure with apo *CjNikZ*, we hypothesized (Hyp.1) that the *Cc*NikZ-II’s K62 was needed for Ni(II) binding. Initial alanine scanning mutagenesis failed to support this, so we assessed holo *CjNikZ*’s binding site with nickel present and found a water network that its E24 was protruding directly into. We hypothesized (Hyp.2) that E60 of *Cc*NikZ-II was therefore critical for Ni(II) binding, which another round of mutagenesis confirmed and provided further evidence to stipulate (Hyp.3) the requirement of a histidine immediately adjacent to this glutamate for high-affinity binding to nickel. To test Hyp.3, we assessed a global phylogeny comprised of NiBPs and PepBPs, then identified eight homologues in addition to *Cc*NikZ-II and *CjNikZ* with diverse v-loop sequences. We found modest conservation of the glutamate and histidine at the front of the core v-loop across several homologues, so we additionally hypothesized (Hyp.4) the core v-loops were modular with respect to their impact on Ni(II) binding affinity. To determine which 6-residue core v-loop sequences to substitute *Cc*NikZ-II’s TEDKYT core v-loop, we used AlphaFold2 predicted homologue structures to generate cognate engineered variants (*i.e*., *Cc*NikZ-II variants) with substituted core v-loops. We found the wild-type NiBPs with the highest Ni(II) binding affinities possessed glutamate and histidine at the front of the core v-loop, thus confirming Hyp.3. We also found the top five wild-type NiBPs’ cognate engineered variants members also had the highest Ni(II) binding affinities, thus confirming Hyp.4. However, the order of the rankings did not match, which we believed to be due to a combination of differences in the v-loop composition and the binding site. This prompted Hyp.5 where we hypothesized the v-loops were modular with respect to their impact on metal promiscuity. Through an automated screening workflow that we developed, we found the tight-binders’ nM-hit profiles could only partially transfer specificity, but this partiality proved useful as it improved nickel specificity relative to the cognate homologue. In the assessment of the μM-hit profiles for low-affinity metals, which was more sporadic, we hypothesized (Hyp.6) that the second coordination sphere and protein matrix were responsible for the wild-type NiBPs’ abilities to discern between low affinity metals. We found the μM-hit profiles for low-affinity metals with the engineered variants (bearing the same *Cc*NikZ-II template) were more likely to be the exact or nearly the same compared to the wild-type NiBPs (bearing different protein matrices), lending additional evidence towards Hyp.6 among other prior observations in this study (*i.e*., mismatched ranking of tight-binders and partial transfer of nM-hit profiles for tight-binders).

As we have demonstrated in this study, NiBPs like *Cc*NikZ-II and its homologues constitute excellent engineering targets given their facile expression and purification, flexible nature (*i.e*., tuneable affinities and specificities), and automatable characterization by liquid-handling workflows. While the literature describing this family of NiBPs has stagnated since the publishing of Lebrette *et al*. (2014), we anticipate more traction in this area given the rise in metalloprotein engineering towards novel bioprocesses to remediate and recover metals from aqueous mining and industrial streams (21, 30, 55, 56). Nickel, cobalt, and REEs like those tested in our metal promiscuity screen constitute strategic metals for a transition to cleaner economies to address climate change.

## Materials and Methods

### Gene cloning and mutagenesis

The open-reading frames encoding for *Cc*NikZ-II and other selected NBPs (Table S3,S4) were synthesized (Twist Bioscience, San Francisco, USA) without the start codon and without their native signal peptides, which were identified using SignalP 5.0 (57). The open-reading frames (codon-optimized for expression in *E. coli*) were cloned into the expression vector p15TV-L (AddGene ID: 26093) under the T7 promoter and in-frame with the N-terminal 6xHisTag (Twist Bioscience). Completed plasmids were transformed into the *E. coli* expression strain LOBSTR (Kerafast, #EC1001) to minimize purification of endogenous Ni(II) binding proteins, and then plated onto LB-agar (carbenicillin, 100 μg/mL). Colonies were inoculated into liquid LB, and the plasmids were miniprepped (GeneAid, #PD300) from overnight inoculants into MilliQ water and sequence verified at the ACGT Sequencing Facility (Toronto, Canada). Glycerol stocks were prepared for storage at −80 °C. All Ala substitution mutant proteins and engineered variants were generated through site-directed mutagenesis (Ranomics, Toronto, Canada) with the TCG codon used for alanine substitution. Sequences were independently verified as described above.

### Protein expression and purification

This protocol is based on prior work and was used to produce all proteins reported in this study (37). Starter cultures were grown from glycerol stock in LB (carbenicillin, 100 μg/mL) for 16 h overnight at 37 °C with shaking. Expression cultures were started by pre-warming TB media (carbenicillin, 100 μg/mL) to 37 °C before 5% v/v inoculation with the starter culture, then grown for 6 hr at 37 °C, then addition of IPTG (BioShop, #IPT002) to 0.4 mM for induction. The expression cultures were then transferred to 16 °C and grown for 16 hr overnight with shaking, then pelleted with centrifugation and transferred to conical vials for one overnight freeze-thaw cycle at −20 °C. Frozen cell pellets were thawed and resuspended in binding buffer (10 mM HEPES, 500 mM NaCl, 5 mM imidazole, pH 7.2) to a final volume of 50-100 mL, followed by addition of 0.25 g lysozyme (BioShop, #LYS702). Cell pellet mixtures were sonicated for 15 min (Q700 Sonicator, Qsonica) and clarified by centrifugation. The soluble layer (supernatant) was applied to a cobalt-charged (Co-)NTA resin (Thermo Fisher, #88221) pre-equilibrated with binding buffer in a gravity-column set-up. Bound proteins were cleansed with wash buffer (10 mM HEPES, 500 mM NaCl, 25 mM imidazole, pH 7.2) and collected with elution buffer (10 mM HEPES, 500 mM NaCl, 250 mM imidazole, pH 7.2). Protein concentrations were determined by Bradford assay, and protein purity was determined by SDS-PAGE analysis (Figure S8) and densitometry on Image Lab 6.0 (Bio-Rad).

Eluted protein was combined with in-house purified TEV protease (1:100 protein-to-TEV) an DTT to 1 mM (BioShop, #TCE101), then transferred to a 10 kDa MWCO dialysis bag (Thermo Fisher, #68100) for dialysis in 4 L dialysis buffer (10 mM HEPES, 1 mM DTT, 1 g/L Chelex 100, pH 7.2) at 4 °C with gentle stirring for 24 hr. Dialyzed samples were then applied to a cobalt-charged NTA resin twice and transferred to a 10 kDa MWCO dialysis bag for dialysis in 4 L activity buffer (AB, 10 mM HEPES, pH 7.2) at 4 °C with gentle stirring for 24 hr and transferred to fresh AB for another 24 hr. Purity was re-confirmed by SDS-PAGE analysis. Finally, they were flash-frozen by liquid nitrogen for storage at −80 °C.

### Protein crystallization and x-ray crystallography

The *apo Cc*NikZ-II crystal was grown at room temperature using the vapor diffusion sitting drop method solution containing 30 mg/mL protein and the reservoir solution 0.1 M HEPES-K buffer (pH 7.5), 30% (w/v) polyethylene glycol 1000. The crystal was cryoprotected using paratone oil. Diffraction data at 100K were collected at a home source Rigaku Micromax 007-HF/R-Axis IV system and data was processed using HKL3000 (58). The structure was solved by Molecular Replacement, using the structure of *apo Cj*NikZ (PDB 4OET)(21) and the CCP4 Mr. Bump program (59). Model building and refinement were performed using Phenix.refine (60) and Coot (61). Geometry was validated using the wwPDB validation server. Atomic coordinates were deposited in the PDB with accession code 8EFZ.

### Phylogenetic and structural analyses

Under the broad family IPR039424 (solute binding protein family 5) in the InterPro database, amino acid sequences from family IPR030678 (peptide/Ni(II) binding protein, MppA-type) were extracted into a FASTA file on April 27^th^ 2019, which contained approximately 150,000 members (43). Sequence redundancy was reduced through the CD-HIT algorithm using an identity cut-off of 0.6 (60%) and default settings (42). Amino acid sequences of *Ec*NikA, *Sa*CntA, *Yp*YntA, *Sa*NikA, *Vp*NikA, *Hh*NikA, *Cj*NikZ, and *Cc*NikZ-II were then manually added (if not already present) to the newly reduced FASTA file, which then had 4,561 members (Supplementary Data File 1). An alignment was performed using the MAFFT v7 algorithm through Geneious v8.1.9 with default settings, and a phylogenetic tree was then built using FastTree v2.1.11 with the Gamma 20 optimization setting (Figure 1, Supplementary Data File 2) (62). Site 1 and Site 2 were extracted from the above tree and imported into JalView v.2.11.2.2 for closer analysis. Sequences of interest (Figure 3) were further extracted and re-aligned again with MAFFT v7. A phylogenetic tree (Figure 3A) was built based on and default settings to examine binding motif conservation.

PyMOL v.2.5.2. was used to visualize protein structures. *Cc*NikZ-II’s structure was predicted on SWISS-MODEL using the apo *Cj*NikZ structure (PDB 4OET) as the homology template (63). It was also predicted using AlphaFold2 through the ColabFold server with the amber settings, and without a template (38, 39). Structural superimposition was performed using built-in PyMOL functions, and rmsd values (Å) were automatically computed (Figure 4). These AlphaFold2 prediction and PyMOL procedures used for *Cc*NikZ-II were applied to all members of the engineered variant set to predict and compare their apo structures with apo *Cc*NikZ-II’s structure reported in this study (Figure 6). Electrostatic surface representations were generated using the APBS biomolecular solvation software suite (64).

### Thermoshift assay

Protein thermostability (*T*_m_, melting temperature) was determined using a SYPRO Orange dye kit (ThermoFisher) with the BioRad CFX96 Real-time PCR Detection System. The FRET (fluorescence resonance energy transfer) channel was set to λ_ex_ 492 nm and λ_em_ 610 nm. Samples were prepared as single replicates using 5 μg protein (generally 1 μL) in a total reaction volume of 20 μL. After 5 min equilibration at 4 °C, the samples were heated from 25-95 °C at a rate of 1 °C per 30 s. Protein unfolding was monitored by changes in the fluorescence of SYPRO orange, and the inflection points from the derivative curves were used to determine the *T*_m_.

### Metal solution preparation and calibration by ICP-MS

All metal solutions were prepared and stored in metal-free tubes (VWR, #89049-172/-176). Metal stock solutions were prepared from chloride salts (BioShop) in non-buffered deionized MilliQ water. The concentrations were then verified by inductively coupled plasma mass spectrometry (ICP-MS) using a ThermoScientific™ iCAP Q ICP-MS system in kinetic energy discrimination (KED) mode. ICP-MS samples of the metals were prepared by dilution into a 2% HNO_3_ solution made from deionized MilliQ water and ultrapure HNO3 Optima™ (Fisher Chemical, #S020101TFIF01). Metal calibration curves were created using a rare earth element standard (Inorganic Ventures, #CCS-1) and a general multi-metal standard (Inorganic Ventures, #IV-28). 0, 100, 500, and 1000 ppb metal standard solutions were prepared with glass volumetric flasks and pipettes previously triple-washed with 10% HNO3 TraceMetal™ (Fisher Chemical, #S010101TFIQ03). Working metal solutions were then made for experiments from these stock solutions using AB prepared from deionized MilliQ water adjusted to the desired pH using NaOH, then filter sterilized by syringe using a 0.2 μm PTFE membrane disc (PALL, #4187). The pH of the working solutions were unchanged based on visual inspection by litmus paper, which was expected based on OLI Studio 9.6 simulations. >99% of the metals were in the aqueous phase based on OLI calculations for all metal concentrations used.

### Metal-affinity determination by microITFQ-LTA

The microplate-based intrinsic tryptophan fluorescence quenching (or ‘ITFQ assays’ for brevity) is based on prior work and was used to characterize all proteins reported in this study (37). ITFQ assays were performed in black, opaque 96-well microplates (Greiner Bio-One, #655076) using the Infinite® M200 (Tecan) plate reader with the settings: fluorescence top-read, 25 °C, λ_ex_ = 280 nm, λ_em_ = 380 nm, and a manual gain of 100. The general procedure used to obtain binding curves by direct titration first required 150 μL of 0.4 μM *Cc*NikZ-II in AB to be added to the appropriate number of wells and equilibrated for 20 min with shaking (orbital, 3mm) to 25 °C. The baseline fluorescence was monitored to ensure equilibration, from which one endpoint reading was made as the blank (*F*_0_). Titrations of metal working solution were incrementally added. Specifically, at each titration increment, a small aliquot (1 to 20 μL) was added by multi-channel pipette, then equilibrated for 3 min with shaking (orbital, 3mm). For larger aliquots (10 to 20 μL), the samples were equilibrated for at least 5 min with shaking (orbital, 3mm), or until the fluorescence stabilized. At each titration step, after the 3-5 min equilibration period, the fluorescence was measured (*F*). This titration procedure was repeated until saturation of the fluorescence signal was observed.

Mixing by pipette was avoided to prevent the introduction of air bubbles or the unintentional removal of protein due to droplets sticking to the inside of pipette tips. To account for any dilution effects, we included a triplicate negative control where the protein was titrated with AB (no added metal) using the same volumes added at each titration step. The inner-filter effect was not observed for the concentrations of metal working solutions used.

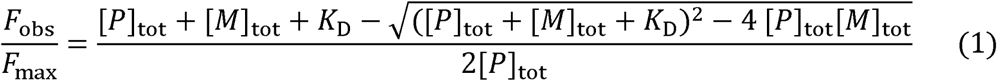

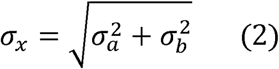

The apparent dissociation constant (*K*_D_) and the maximum fluorescence change (*F*_max_) were determined by fitting a 1:1 stoichiometry mass-action kinetic model (Eqn.1) to the data (Table S2) (34). Here, [*M*]_tot_ and [*P*]_tot_ represent the total metal and protein concentration in the well at each titration step. The observed fluorescence change *F*_obs_ is calculated by subtracting the fluorescence of the sample when no metal is added (*F*_0_) from the fluorescence at each titration step where metal is added (*F*). Experiments were performed in triplicates. The standard deviation of the resulting *F*_obs_ is therefore calculated by error propagation (Eqn.2). The standard deviations of *K*_D_ and *F*_max_ are computed by the scipy.optimize function in Python where this data was analyzed (Supplementary Data File 4, 5) (65). Plots were generated using matplotlib in Python (66).

### Automated screening of purified proteins for metal binding

A modified ITFQ assay was used to detect changes in fluorescence upon metal binding (37). 150 μL of 0.4 μM *Cc*NikZ-II in AB was added to wells (Greiner Bio-One, #655076). Specifically, for each purified protein triplicates (three wells) were prepared for titrating 11 metals (Ni^2+^, Co^2+^, Cu^2+^, Zn^2+^, Mn^2+^, Al^3+^, Pb^2+^, Cd^2+^, La^3+^, Eu^3+^, and Yb^3+^ as chloride salts) at final metal concentrations of 0, 0.1, and 100 μM each. The handling of this liquid and the titration of the metals from a 96-well deep block (VWR, #82051-260), was performed by a EvoWare v2.7 scripted Tecan Freedom Evo 100 base equipped with a Tecan fixed-tip liquid handling arm, a Tecan RoMa (robotic manipulator) arm, and a Tecan Infinite M200 plate reader (same as one used in Section 2.7). Fluorescence readings were taken after equilibration for 20 min with shaking (orbital, 3mm) at 25 °C to obtain the blank *F*_0_, and after allowing for 5 min equilibration with shaking (orbital, 3mm) upon titrating the metals to obtain *F*. The fold-changes in fluorescence signal between the 100 nM and 100 μM titrations (*F*) each compared to the negative control blank (*F*_0_) were calculated and plotted using matplotlib in Python (66).

---

## Acknowledgements

The authors thank…

## Author contributions

PD performed…

## Abbreviations

–

## References

1. Begg, S. L. (2019) The role of metal ions in the virulence and viability of bacterial pathogens. Biochemical Society Transactions. 47, 77–87

2. Chandrangsu, P., Rensing, C., and Helmann, J. D. (2017) Metal homeostasis and resistance in bacteria. Nat Rev Microbiol. 15, 338–350

3. Haferburg, G., and Kothe, E. (2007) Microbes and metals: interactions in the environment. Journal of Basic Microbiology. 47, 453–467

4. Foster, A. W., Osman, D., and Robinson, N. J. (2014) Metal Preferences and Metallation. Journal of Biological Chemistry. 289, 28095–28103

5. Dudev, T., and Lim, C. (2014) Competition among Metal Ions for Protein Binding Sites: Determinants of Metal Ion Selectivity in Proteins. Chem. Rev. 114, 538–556

6. Barwinska-Sendra, A., and Waldron, K. J. (2017) The Role of Intermetal Competition and Mis-Metalation in Metal Toxicity. in Advances in Microbial Physiology, pp. 315–379, Elsevier, 70, 315–379

7. Osman, D., Martini, M. A., Foster, A. W., Chen, J., Scott, A. J. P., Morton, R. J., Steed, J. W., Lurie-Luke, E., Huggins, T. G., Lawrence, A. D., Deery, E., Warren, M. J., Chivers, P. T., and Robinson, N. J. (2019) Bacterial sensors define intracellular free energies for correct enzyme metalation. Nat Chem Biol. 15, 241–249

8. Chandravanshi, M., Tripathi, S. K., and Kanaujia, S. P. (2021) An updated classification and mechanistic insights into ligand binding of the substrate-binding proteins. FEBS Letters. 595, 2395–2409

9. Scheepers, G. H., Lycklama a Nijeholt, J. A., and Poolman, B. (2016) An updated structural classification of substrate-binding proteins. FEBS Letters. 590, 4393–4401

10. Berntsson, R. P.-A., Smits, S. H. J., Schmitt, L., Slotboom, D.-J., and Poolman, B. (2010) A structural classification of substrate-binding proteins. FEBS Letters. 584, 2606–2617

11. Tam, R., and Saier, M. H. (1993) Structural, functional, and evolutionary relationships among extracellular solute-binding receptors of bacteria. Microbiol Rev. 57, 320–346

12. Navarro, C., Wu, L.-F., and Mandrand-Berthelot, M.-A. (1993) The nik operon of Escherichia coli encodes a periplasmic binding-protein-dependent transport system for nickel. Molecular Microbiology. 9, 1181–1191

13. De Pina, K., Navarro, C., Mcwalter, L., Boxer, D. H., Price, N. C., Kelly, S. M., Mandrand-Berthelot, M.-A., and Wu, L.-F. (1995) Purification and Characterization of the Periplasmic Nickel-Binding Protein NikA of Escherichia coli K12. European Journal of Biochemistry. 227, 857–865

14. Heddle, J., Scott, D. J., Unzai, S., Park, S.-Y., and Tame, J. R. H. (2003) Crystal Structures of the Liganded and Unliganded Nickel-binding Protein NikA from Escherichia coli*. Journal of Biological Chemistry. 278, 50322–50329

15. Cherrier, M. V., Martin, L., Cavazza, C., Jacquamet, L., Lemaire, D., Gaillard, J., and Fontecilla-Camps, J. C. (2005) Crystallographic and Spectroscopic Evidence for High Affinity Binding of FeEDTA(H2O)- to the Periplasmic Nickel Transporter NikA. J. Am. Chem. Soc. 127, 10075–10082

16. Rowe, J. L., Starnes, G. L., and Chivers, P. T. (2005) Complex Transcriptional Control Links NikABCDE-Dependent Nickel Transport with Hydrogenase Expression in Escherichia coli. Journal of Bacteriology. 187, 6317–6323

17. Cherrier, M. V., Cavazza, C., Bochot, C., Lemaire, D., and Fontecilla-Camps, J. C. (2008) Structural Characterization of a Putative Endogenous Metal Chelator in the Periplasmic Nickel Transporter NikA. Biochemistry. 47, 9937–9943

18. Cherrier, M. V., Girgenti, E., Amara, P., Iannello, M., Marchi-Delapierre, C., Fontecilla-Camps, J. C., Ménage, S., and Cavazza, C. (2012) The structure of the periplasmic nickel-binding protein NikA provides insights for artificial metalloenzyme design. J Biol Inorg Chem. 17, 817–829

19. Lebrette, H., Iannello, M., Fontecilla-Camps, J. C., and Cavazza, C. (2013) The binding mode of Ni-(L-His)2 in NikA revealed by X-ray crystallography. Journal of Inorganic Biochemistry. 121, 16–18

20. Howlett, R. M., Hughes, B. M., Hitchcock, A., and Kelly, D. J. (2012) Hydrogenase activity in the foodborne pathogen Campylobacter jejuni depends upon a novel ABC-type nickel transporter (NikZYXWV) and is SlyD-independent. Microbiology. 158, 1645–1655

21. Lebrette, H., Brochier-armanet, C., Zambelli, B., de Reuse, H., Borezée-Durant, E., Ciurli, S., and Cavazza, C. (2014) Promiscuous Nickel Import in Human Pathogens: Structure, Thermodynamics, and Evolution of Extracytoplasmic Nickel-Binding Proteins. Structure. 22, 1421–1432

22. Lebrette, H., Borezée-Durant, E., Martin, L., Richaud, P., Boeri Erba, E., and Cavazza, C. (2015) Novel insights into nickel import in Staphylococcus aureus: the positive role of free histidine and structural characterization of a new thiazolidine-type nickel chelator†. Metallomics. 7, 613–621

23. Benoit, S. L., Seshadri, S., Lamichhane-Khadka, R., and Maier, R. J. (2013) Helicobacter hepaticus NikR controls urease and hydrogenase activities via the NikABDE and HH0418 putative nickel import proteins. Microbiology. 159, 136–146

24. Shaik, M. M., Cendron, L., Salamina, M., Ruzzene, M., and Zanotti, G. (2014) Helicobacter pylori periplasmic receptor CeuE (HP1561) modulates its nickel affinity via organic metallophores. Molecular Microbiology. 91, 724–735

25. Park, K.-S., Iida, T., Yamaichi, Y., Oyagi, T., Yamamoto, K., and Honda, T. (2000) Genetic Characterization of DNA Region Containing the trh and ure Genes of Vibrio parahaemolyticus. Infection and Immunity. 68, 5742–5748

26. Hughes, A., Wilson, S., Dodson, E. J., Turkenburg, J. P., and Wilkinson, A. J. (2019) Crystal structure of the putative peptide-binding protein AppA from Clostridium difficile. Acta Cryst F. 75, 246–253

27. Wegner, S. V., Boyaci, H., Chen, H., Jensen, M. P., and He, C. (2009) Engineering A Uranyl-Specific Binding Protein from NikR. Angewandte Chemie International Edition. 48, 2339–2341

28. Jia, X., Ma, Y., Bu, R., Zhao, T., and Wu, K. (2020) Directed evolution of a transcription factor PbrR to improve lead selectivity and reduce zinc interference through dual selection. AMB Express. 10, 67

29. Zhou, L., Bosscher, M., Zhang, C., Özçubukçu, S., Zhang, L., Zhang, W., Li, C. J., Liu, J., Jensen, M. P., Lai, L., and He, C. (2014) A protein engineered to bind uranyl selectively and with femtomolar affinity. Nature Chem. 6, 236–241

30. Dong, Z., Mattocks, J. A., Deblonde, G. J.-P., Hu, D., Jiao, Y., Cotruvo, J. A., and Park, D. M. (2021) Bridging Hydrometallurgy and Biochemistry: A Protein-Based Process for Recovery and Separation of Rare Earth Elements. ACS Cent. Sci. 7, 1798–1808

31. Keshav, V., Franklyn, P., and Kondiah, K. (2019) Recombinant Fusion Protein PbrD Cross-Linked to Calcium Alginate Nanoparticles for Pb Remediation. ACS Omega. 4, 16816–16825

32. Garner, K. L. (2021) Principles of synthetic biology. Essays in Biochemistry. 65, 791–811

33. Carbonell, P., Jervis, A. J., Robinson, C. J., Yan, C., Dunstan, M., Swainston, N., Vinaixa, M., Hollywood, K. A., Currin, A., Rattray, N. J. W., Taylor, S., Spiess, R., Sung, R., Williams, A. R., Fellows, D., Stanford, N. J., Mulherin, P., Le Feuvre, R., Barran, P., Goodacre, R., Turner, N. J., Goble, C., Chen, G. G., Kell, D. B., Micklefield, J., Breitling, R., Takano, E., Faulon, J.-L., and Scrutton, N. S. (2018) An automated Design-Build-Test-Learn pipeline for enhanced microbial production of fine chemicals. Commun Biol. 1, 1–10

34. Weert, M. van de, and Stella, L. (2011) Fluorescence quenching and ligand binding: A critical discussion of a popular methodology. Journal of Molecular Structure. 998, 144–150

35. Callis, P. R. (2014) Binding phenomena and fluorescence quenching. I: Descriptive quantum principles of fluorescence quenching using a supermolecule approach. Journal of Molecular Structure. 1077, 14–21

36. Callis, P. R. (2014) Binding phenomena and fluorescence quenching. II: Photophysics of aromatic residues and dependence of fluorescence spectra on protein conformation. Journal of Molecular Structure. 1077, 22–29

37. Diep, P., Mahadevan, R., and Yakunin, A. F. (2020) A microplate screen to estimate metal-binding affinities of metalloproteins. Analytical Biochemistry. 609, 113836

38. Jumper, J., Evans, R., Pritzel, A., Green, T., Figurnov, M., Ronneberger, O., Tunyasuvunakool, K., Bates, R., Žídek, A., Potapenko, A., Bridgland, A., Meyer, C., Kohl, S. A. A., Ballard, A. J., Cowie, A., Romera-Paredes, B., Nikolov, S., Jain, R., Adler, J., Back, T., Petersen, S., Reiman, D., Clancy, E., Zielinski, M., Steinegger, M., Pacholska, M., Berghammer, T., Bodenstein, S., Silver, D., Vinyals, O., Senior, A. W., Kavukcuoglu, K., Kohli, P., and Hassabis, D. (2021) Highly accurate protein structure prediction with AlphaFold. Nature. 596, 583–589

39. Mirdita, M., Schütze, K., Moriwaki, Y., Heo, L., Ovchinnikov, S., and Steinegger, M. (2022) ColabFold: making protein folding accessible to all. Nat Methods. 19, 679–682

40. Holm, L., and Rosenstrï¿½m, P. (2010) Dali server: conservation mapping in 3D. Nucleic Acids Research. 38, W545–W549

41. Diep, P., Mahadevan, R., and Yakunin, A. (2019) A microplate screen for metal-binding activity based on a nickel-binding protein from Clostridium carboxidivorans. bioRxiv. 10.1101/820670

42. Li, W., and Godzik, A. (2006) Cd-hit: a fast program for clustering and comparing large sets of protein or nucleotide sequences. Bioinformatics. 22, 1658–1659

43. Blum, M., Chang, H.-Y., Chuguransky, S., Grego, T., Kandasaamy, S., Mitchell, A., Nuka, G., Paysan-Lafosse, T., Qureshi, M., Raj, S., Richardson, L., Salazar, G. A., Williams, L., Bork, P., Bridge, A., Gough, J., Haft, D. H., Letunic, I., Marchler-Bauer, A., Mi, H., Natale, D. A., Necci, M., Orengo, C. A., Pandurangan, A. P., Rivoire, C., Sigrist, C. J. A., Sillitoe, I., Thanki, N., Thomas, P. D., Tosatto, S. C. E., Wu, C. H., Bateman, A., and Finn, R. D. (2021) The InterPro protein families and domains database: 20 years on. Nucleic Acids Research. 49, D344–D354

44. Price, M. N., Dehal, P. S., and Arkin, A. P. (2010) FastTree 2 – Approximately Maximum-Likelihood Trees for Large Alignments. PLOS ONE. 5, e9490

45. Young, T. R., and Xiao, Z. (2021) Principles and practice of determining metal–protein affinities. Biochemical Journal. 478, 1085–1116

46. Li, W.-J., Kroppenstedt, R. M., Wang, D., Tang, S.-K., Lee, J.-C., Park, D.-J., Kim, C.-J., Xu, L.-H., and Jiang, C.-L. 2006 Five novel species of the genus Nocardiopsis isolated from hypersaline soils and emended description of Nocardiopsis salina Li et al. 2004. International Journal of Systematic and Evolutionary Microbiology. 56, 1089–1096

47. Zavarzina, D. G., Sokolova, T. G., Tourova, T. P., Chernyh, N. A., Kostrikina, N. A., and Bonch-Osmolovskaya, E. A. (2007) Thermincola ferriacetica sp. nov., a new anaerobic, thermophilic, facultatively chemolithoautotrophic bacterium capable of dissimilatory Fe(III) reduction. Extremophiles. 11, 1–7

48. Irving, H., and Williams, R. J. P. (2004) 637. The stability of transition-metal complexes. Journal of the Chemical Society (Resumed). 10.1039/jr9530003192

49. Eom, H., and Song, W. J. (2019) Emergence of metal selectivity and promiscuity in metalloenzymes. J Biol Inorg Chem. 24, 517–531

50. Young, T. R., Martini, M. A., Foster, A. W., Glasfeld, A., Osman, D., Morton, R. J., Deery, E., Warren, M. J., and Robinson, N. J. (2021) Calculating metalation in cells reveals CobW acquires CoII for vitamin B12 biosynthesis while related proteins prefer ZnII. Nat Commun. 12, 1195

51. Osman, D., Martini, M. A., Foster, A. W., Chen, J., Scott, A. J. P., Morton, R. J., Steed, J. W., Lurie-Luke, E., Huggins, T. G., Lawrence, A. D., Deery, E., Warren, M. J., Chivers, P. T., and Robinson, N. J. (2019) Bacterial sensors define intracellular free energies for correct enzyme metalation. Nat Chem Biol. 15, 241–249

52. Neville, S. L., Sjöhamn, J., Watts, J. A., MacDermott-Opeskin, H., Fairweather, S. J., Ganio, K., Carey Hulyer, A., McGrath, A. P., Hayes, A. J., Malcolm, T. R., Davies, M. R., Nomura, N., Iwata, S., O’Mara, M. L., Maher, M. J., and McDevitt, C. A. (2021) The structural basis of bacterial manganese import. Sci Adv. 7, eabg3980

53. Lawrence, M. C., Pilling, P. A., Epa, V. C., Berry, A. M., Ogunniyi, A. D., and Paton, J. C. (1998) The crystal structure of pneumococcal surface antigen PsaA reveals a metal-binding site and a novel structure for a putative ABC-type binding protein. Structure. 6, 1553–1561

54. Heddle, J., Scott, D. J., Unzai, S., Park, S.-Y., and Tame, J. R. H. (2003) Crystal Structures of the Liganded and Unliganded Nickel-binding Protein NikA from Escherichia coli*. Journal of Biological Chemistry. 278, 50322–50329

55. Capeness, M. J., and Horsfall, L. E. (2020) Synthetic biology approaches towards the recycling of metals from the environment. Biochemical Society Transactions. 48, 1367–1378

56. Urbina, J., Patil, A., Fujishima, K., Paulino-Lima, I. G., Saltikov, C., and Rothschild, L. J. (2019) A new approach to biomining: Bioengineering surfaces for metal recovery from aqueous solutions. Sci Rep. 9, 16422

57. Almagro Armenteros, J. J., Tsirigos, K. D., Sønderby, C. K., Petersen, T. N., Winther, O., Brunak, S., von Heijne, G., and Nielsen, H. (2019) SignalP 5.0 improves signal peptide predictions using deep neural networks. Nat Biotechnol. 37, 420–423

58. Minor, W., Cymborowski, M., Otwinowski, Z., and Chruszcz, M. (2006) HKL-3000: the integration of data reduction and structure solution – from diffraction images to an initial model in minutes. Acta Cryst D. 62, 859–866

59. Keegan, R. M., McNicholas, S. J., Thomas, J. M. H., Simpkin, A. J., Simkovic, F., Uski, V., Ballard, C. C., Winn, M. D., Wilson, K. S., and Rigden, D. J. (2018) Recent developments in MrBUMP: better search-model preparation, graphical interaction with search models, and solution improvement and assessment. Acta Cryst D. 74, 167–182

60. Liebschner, D., Afonine, P. V., Baker, M. L., Bunkóczi, G., Chen, V. B., Croll, T. I., Hintze, B., Hung, L.-W., Jain, S., McCoy, A. J., Moriarty, N. W., Oeffner, R. D., Poon, B. K., Prisant, M. G., Read, R. J., Richardson, J. S., Richardson, D. C., Sammito, M. D., Sobolev, O. V., Stockwell, D. H., Terwilliger, T. C., Urzhumtsev, A. G., Videau, L. L., Williams, C. J., and Adams, P. D. (2019) Macromolecular structure determination using X-rays, neutrons and electrons: recent developments in Phenix. Acta Cryst D. 75, 861–877

61. Emsley, P., Lohkamp, B., Scott, W. G., and Cowtan, K. (2010) Features and development of Coot. Acta Cryst D. 66, 486–501

62. Katoh, K., Misawa, K., Kuma, K., and Miyata, T. (2002) MAFFT: a novel method for rapid multiple sequence alignment based on fast Fourier transform. Nucleic Acids Research. 30, 3059–3066

63. Waterhouse, A., Bertoni, M., Bienert, S., Studer, G., Tauriello, G., Gumienny, R., Heer, F. T., de Beer, T. A. P., Rempfer, C., Bordoli, L., Lepore, R., and Schwede, T. (2018) SWISS-MODEL: homology modelling of protein structures and complexes. Nucleic Acids Research. 46, W296–W303

64. Jurrus, E., Engel, D., Star, K., Monson, K., Brandi, J., Felberg, L. E., Brookes, D. H., Wilson, L., Chen, J., Liles, K., Chun, M., Li, P., Gohara, D. W., Dolinsky, T., Konecny, R., Koes, D. R., Nielsen, J. E., Head-Gordon, T., Geng, W., Krasny, R., Wei, G.-W., Holst, M. J., McCammon, J. A., and Baker, N. A. (2018) Improvements to the APBS biomolecular solvation software suite. Protein Sci. 27, 112–128

65. Virtanen, P., Gommers, R., Oliphant, T. E., Haberland, M., Reddy, T., Cournapeau, D., Burovski, E., Peterson, P., Weckesser, W., Bright, J., van der Walt, S. J., Brett, M., Wilson, J., Millman, K. J., Mayorov, N., Nelson, A. R. J., Jones, E., Kern, R., Larson, E., Carey, C. J., Polat, İ., Feng, Y., Moore, E. W., VanderPlas, J., Laxalde, D., Perktold, J., Cimrman, R., Henriksen, I., Quintero, E. A., Harris, C. R., Archibald, A. M., Ribeiro, A. H., Pedregosa, F., van Mulbregt, P., SciPy 1.0 Contributors, Vijaykumar, A., Bardelli, A. P., Rothberg, A., Hilboll, A., Kloeckner, A., Scopatz, A., Lee, A., Rokem, A., Woods, C. N., Fulton, C., Masson, C., Häggström, C., Fitzgerald, C., Nicholson, D. A., Hagen, D. R., Pasechnik, D. V., Olivetti, E., Martin, E., Wieser, E., Silva, F., Lenders, F., Wilhelm, F., Young, G., Price, G. A., Ingold, G.-L., Allen, G. E., Lee, G. R., Audren, H., Probst, I., Dietrich, J. P., Silterra, J., Webber, J. T., Slavič, J., Nothman, J., Buchner, J., Kulick, J., Schönberger, J. L., de Miranda Cardoso, J. V., Reimer, J., Harrington, J., Rodríguez, J. L. C., Nunez-Iglesias, J., Kuczynski, J., Tritz, K., Thoma, M., Newville, M., Kümmerer, M., Bolingbroke, M., Tartre, M., Pak, M., Smith, N. J., Nowaczyk, N., Shebanov, N., Pavlyk, O., Brodtkorb, P. A., Lee, P., McGibbon, R. T., Feldbauer, R., Lewis, S., Tygier, S., Sievert, S., Vigna, S., Peterson, S., More, S., Pudlik, T., Oshima, T., Pingel, T. J., Robitaille, T. P., Spura, T., Jones, T. R., Cera, T., Leslie, T., Zito, T., Krauss, T., Upadhyay, U., Halchenko, Y. O., and Vázquez-Baeza, Y. (2020) SciPy 1.0: fundamental algorithms for scientific computing in Python. Nat Methods. 17, 261–272

66. Hunter, J. D. (2007) Matplotlib: A 2D Graphics Environment. Computing in Science & Engineering. 9, 90–95

