## Supplementary material for "Ni(II) binding affinity and specificity of solute binding proteins: the importance of the double His motif and variable loop revealed by structural and mutational studies": SI

*From the ^1^Department of Chemical Engineering and Applied Chemistry, BioZone – Centre for Applied Bioscience and Bioengineering, University of Toronto, Toronto, Ontario, Canada; ^2^Department of Microbiology, Immunology and Infectious Diseases, University of Calgary, Calgary, Alberta, Canada; ^3^Institute of Biomedical Engineering, University of Toronto, Toronto, Ontario, Canada; ^4^Centre for Environmental Biotechnology, School of Natural Sciences, Bangor University, Bangor, Wales, United Kingdom*

**Supplementary Tables**

Table S1. X-ray crystallographic statistics for the structure of *Cc*NikZ-II, (p2)

Table S2. Nickel binding parameters of the wild type and engineered NiBPs, (p3)

Table S3. NiBPs characterized in this study, (p4)

Table S4. Amino acid sequences of *Cc*NikZ-II and other NiBPs characterized in this study, (p5)

**Supplementary Figures**

Figure S1. Binding curves of mutant library members, (p6)

Figure S2. Full Multiple Sequence Alignment, (p7-8)

Figure S3. Comparison of (A) SWISS-MODEL and (B) AlphaFold 2 predicted structures of apo CcNikZ-II structures with crystal structure, (p9)

Figure S4. Binding curves of homologue library members, (p10)

Figure S5. Binding curves of exchange library members, (p11)

Figure S6. Log-binding curves of library members with the lowest K_D_ values (highest affinity) , (p12)

Figure S7. Melting point determination by thermal shift assay, (p13)

Figure S8. SDS-PAGE analysis of purified NiBP proteins used in this study, (p14)

**Table S1.** X-ray crystallographic statistics for the structure of apo CcNikZ-II.

| ***Data collection*** |  |
| --- | --- |
| Space group | P2 |
| Unit cell  *a*, *b, c* (Å)  α, β, γ, (°) | 73.6, 110.9, 73.7  90, 113.7, 90 |
| Resolution, Å | 30.00 – 2.38 |
| R*_merge_*^a^  R*_pim_*^b^ | 0.101 (0.536)*  0.057 (0.267) |
| CC_1/2_ | 0.990 (0.652) |
| *I* / σ(*I)* | 20.66 (1.64) |
| Completeness, % | 93.9 (81.6) |
| Redundancy | 4.2 (3.4) |
| ***Refinement*** |  |
| Resolution, Å | 29.71 – 2.38 |
| No. unique reflections:  working, test | 39058, 2057 |
| *R*-factor/free *R­*-factor^c^ | 22.8/25.9 (33.3/36.4) |
| No. refined atoms, molecules  Protein  Solvent  Water | 7781, 2  2  172 |
| *B*-factors  Protein  Solvent  Water | 51.9  42.1  43.4 |
| r.m.s.d.  Bond lengths, Å  Bond angles, ° | 0.002  0.545 |

*values in brackets refer to highest resolution shells.

^a^*R*_merge_ = Σ_hkl_Σ_j_|*I*_hkl.j_ - 〈*I*_hkl_〉|/Σ_hkl_Σ_j_I_hk,j_, where *I*_hkl,j_ and 〈*I*_hkl_〉 are the *j*th and mean measurement of the intensity of reflection *j*.

^b^*R*_pim_ = Σ_hkl_√(n/n-1) Σ^n^_j=1_|*I*_hkl.j_ - 〈*I*_hkl_〉|/Σ_hkl_Σ_j_I_hk,j_

^c^*R* = Σ|F_p_^obs^ – F_p_^calc^|/ΣF_p_^obs^, where F_p_^obs^ and F_p_^calc^ are the observed and calculated structure factor amplitudes, respectively.

**Table S2.** Nickel binding parameters of the wild type and engineered NiBPs

| **NiBPs** | **ID** | ***K*_D_  (μM)** | ***K*_D_  SD** | ***K*_D_  RSD** | ***F*_max_ (a.u.)** | ***F*_max_  SD** | ***F*_max_**  **RSD** |
| --- | --- | --- | --- | --- | --- | --- | --- |
| wild type  nickel-biding  proteins | *Ng*NikZ | 0.06 ^a^ | 0.04 | 67% | 764.95 | 22.35 | 3% |
|  | *Cj*NikZ | 0.15 ^a^ | 0.09 | 60% | 497.93 | 22.21 | 4% |
|  | *Bo*NikZ | 0.22 ^a^ | 0.06 | 27% | 1183.92 | 26.07 | 2% |
|  | *Mp*NikZ | 0.66 ^a^ | 0.18 | 27% | 819.92 | 22.94 | 3% |
|  | *Cf*NikZ | 0.87 ^a^ | 0.29 | 33% | 367.2 | 13.19 | 4% |
|  | *Tf*NikZ | 2.52 | 0.63 | 25% | 483.25 | 18.35 | 4% |
|  | *Cs*NikZ | 2.69 | 1.04 | 39% | 383.44 | 23.26 | 6% |
|  | *Um*NikZ | 4.72 | 1.09 | 23% | 1173.61 | 54.5 | 5% |
|  | *Su*NikZ | 5.91 | 2.01 | 34% | 570.34 | 42.59 | 7% |
|  | *Cc*NikZ-II | 9.97 | 1.83 | 18% | 456.67 | 22.33 | 5% |
|  | *Vb*NikZ ^b^ | - | - | - | - | - | - |
| Ala substitution  mutants | K33A | 2.31 | 0.49 | 21% | 442.6 | 13.83 | 3% |
|  | TEDKYT-AAAHAA | 4.49 | 0.42 | 9% | 338.71 | 6.29 | 2% |
|  | T30A | 5.41 | 0.73 | 13% | 898.74 | 25.67 | 3% |
|  | T35A | 9.05 | 1.89 | 21% | 412.65 | 22.08 | 5% |
|  | Y34A | 10.01 | 1.83 | 18% | 1197.92 | 58.16 | 5% |
|  | NEE-AAA | 10.68 | 1.54 | 14% | 381.44 | 14.93 | 4% |
|  | H483A | 12.15 | 1.82 | 15% | 428.19 | 18.31 | 4% |
|  | D32A | 13.88 | 3.92 | 28% | 247.16 | 20.78 | 8% |
|  | H482A | 21.26 | 3.26 | 15% | 352.59 | 18.6 | 5% |
|  | TEDKYT-AAAAAA | 35.64 | 8.72 | 24% | 232.02 | 23.25 | 10% |
|  | TEDKYT-AAAKAA | 45.09 | 25.34 | 56% | 217.15 | 54.18 | 25% |
|  | E31A | 84.34 | 48.79 | 58% | 538.01 | 171.65 | 32% |
|  | R350A ^b^ | - | - | - | - | - | - |
|  | H482A, H483A ^b^ | - | - | - | - | - | - |
| engineering  *Cc*NikZ-II  variants | TEDKYT-DEDHDT *(Cf>Cc)* | 0.67^a^ | 0.22 | 33% | 345.56 | 11.37 | 3% |
|  | TEDKYT-DEHGEI (*Bo*>*Cc*) | 1.09 ^a^ | 0.14 | 13% | 598.65 | 8.6 | 1% |
|  | TEDKYT-YEHGEI *(Mp*>*Cc )* | 1.98 | 0.27 | 14% | 828.6 | 15.51 | 2% |
|  | TEDKYT-DEHLDP *(Ng*>*Cc )* | 2.1 | 0.48 | 23% | 455.29 | 14.58 | 3% |
|  | TEDKYT-SEDHDA *(Cj*>*Cc)* | 2.68 | 0.8 | 30% | 295.81 | 13.77 | 5% |
|  | TEDKYT-NTHDEL (*Vb*>*Cc*) | 3.07 | 0.36 | 12% | 1029.35 | 20.05 | 2% |
|  | TEDKYT-DESQEI *(Um*>*Cc)* | 4.02 | 0.56 | 14% | 586.51 | 15.32 | 3% |
|  | TEDKYT-EETNLD *(Su*>*Cc)* | 4.04 | 1.02 | 25% | 577.57 | 27.38 | 5% |
|  | TEDKYT-IESHAF *(Cd*>*Cc)* | 4.34 | 1.18 | 27% | 1073.65 | 56.43 | 5% |
|  | TEDKYT-DDSPVD *(Tf*>*Cc )* | 7.63 | 0.81 | 11% | 381.35 | 9.81 | 3% |

^a^ nanomolar affinity variants should be interpreted as upper bounds for *K*_D_, and require competition assays to more accurately determine the true apparent *K*_D_ value
^b^ proteins that could not be expressed using the same protocol as the other proteins reported in this study

**Table S3.** NiBPs characterized in this study

| Protein Name  (this study) | NCBI ID | GenBank | Microorganism |
| --- | --- | --- | --- |
| *Bo*NikZ | WP_019161422.1 |  | *Blautia obeum* |
| *Cc*NikZ-II |  | CP011803.1 | *Clostridium carboxidivorans* |
| *Cj*NikZ | WP_087695781.1 |  | *Campylobacter jejuni* |
| *Cf*NikZ | WP_038454238.1 |  | *Campylobacter fetus* |
| *Cs*NikZ | WP_029161324.1 |  | *Clostridium scatologenes* |
| *Mp*NikZ |  | CP016199.1 | *Mogibacterium pumilum* |
| *Ng*NikZ |  | CP022753.1 | *Nocardiopsis gilva* |
| *Su*NikZ |  | CP015109.1 | *Sporosarcina ureae* |
| *Tf*NikZ | WP_013120112.1 |  | *Thermincola ferriacetica* |
| *Um*NikZ |  | KGR79695.1 | *Ureibacillus manganicus* |
| *Vb*NikZ |  | KXB90791.1 | *Veillonellaceae bacterium* |

**Table S4.** Amino acid sequences of CcNikZ-II and other NiBPs characterized in this study

| *Bo*NikZ | EEETTLVYGSGDYTRINPAMDEHGEINILIFNGLTAHDGDNQVVPGLAESWDFDDETNTYTFHMAEDAKWQDGEPVTAEDVKFTIEAIMDPENGSENAPNYEDVEEINVIDDHTVAFKLEDKNVAFLDYMTMAVLPKHLLEGEDMQTSDFFRNPVGTGPYKIESWDEGQAITLVKNEDYFKGEPSIDKIVFKIVPDDNAKALQLKSGELDLALLTPKDAAAFADDEVYTCYDMKTSDYRGIMFNFGNEYWQKNRDLIPAVCYGLDRQAIIDAVLLGQGMPAYGPLQRNVYNYEDVEHYDYNPEKSKEILEAAGCEMGDDGFYYRDGEKVGFVISVSAGDQVRIDMAQIAAQELKEIGMDVSVEIPAQTDWAGQMAFLIGWGSPFDADDHTYKVFGTDKGANYSGYSNADVDKYLTEARQSADPEVRAEAYANFQKALAEDPAYAMICYIDANYVADSNIKGIDPDTIMGHHGVGIFWNVADWTIE |
| --- | --- |
| *Cc*NikZ-II | TSTKKASDSGKTLVYGAEFEDEKLNPILTEDKYTNEEIFTGLMKFDENNIPKPEIADSYTISDDKLTYDFKLKKGIKFHDETELKAEDVVFTLKSILDEKVNSSLKPEYSEIKDVQAVNDYEVKVILKEAFPPLLDKLTIGIIPKHCFNGKDINTAEFNQKPIGTGPFKFVKWDKGNNITLTKFKDYYGKTGNIEKFVVKFIADYNVRAMQLQTGEIDVAYLEPSQVSKIEKLNNVKVYKVDTADYRCMMYNMKKDIWKDVNVRKAFNYALDRKGMVDGILLGYGSEAYSPLQINKFNNPDVEKYSYNLDKSNSLLESAGWKKGSDGIRVKDGKKLEFTLTAPKTDEVRVKMAEYFASQFKKIGAEVKVDALDWDAIKIDKCDAFLLGWGSPFDADDHTFRLFHSSEINGGDNNGSYSNPKVDEALYKARTTTDENERKKYYAEFQKELAEDPAYDFGVYLKALFGVNKRVSGVKEKVLGHHGAGFLWNVEQWNVN |
| *Cf*NikZ | ATPKDTIVIGIENETSRINPLFDEDHDTALDFVFSGLTKFTLDMKIEPDLAKSWKVSEDGLVWTFYLRDDALWHDGEKFSAEDVKFTIEQALNDKLNAPAKASFEEVQSVEVINKYELKIMLKSPFPPLLDALSIGVLPKHILDGKDINSDKFNQNPIGTGPFKFKKWQKGSYISFEANKDFYRGKPKADKVILKIVPDYNIRTYEIKRGDLDVALIESNLVKELQNDKNIKIMKLDSADYRALMFNFDNEILKSLAVRQAINYAIDRELISTKILHGYGFSANNPIQKSWANDKNAKIYEYNIEKAKEILANDGWELKNKVLQKNGKRLEFDIYAFNGDPFRVTLANIIQSELSKIGIKARAIAKNHGAFEIDKVDSFIIGWGSPLDPDVQTYRVFSSKMDAAKNENGWNYNHYSDKTVDETLLKARTTDKTEERKLWYSKFINALHEDPPFAFLVYLQYPLAYSDKISGIKPTILGHHGARFTHNIEEWSKN |
| *Cj*NikZ | KIPKDTLIIAVENEIARINPAYSEDHDAVINLVFSGLTRFDENMSLKPDLAKSWDISKDGLVYDIFLRDDVLWHDGVKFSADDVKFSIEAFKNPKNNSSIYVNFEDIKSVEILNPSHVKITLFKPYPAFLDALSIGMLPKHLLENENLNTSSFNQNPIGTGPYKFVKWKKGEYVEFKANEHFYLDKVKTPRLIIKHIFDPSIASAELKNGKIDAALIDVSLLNIFKNDENFGILREKSADYRALMFNLDNEFLKDLKVRQALNYAVDKESIVKNLLHDYAFVANHPLERSWANSKNFKIYKYDPKKAEDLLVSAGFKKNKDGNFEKDGKILEFEIWAMSNDPLRVSLAGILQSEFRKIGVVSKVVAKPAGSFDYSKVDSFLIGWGSPLDPDFHTFRVFESSQDSALNDEGWNFGHYHDKKVDIALQKARNTSNLEERKKYYKDFIDALYENPPFIFLAYLDFALVYNKDLKGIKTRTLGHHGVGFTWNVYEWSK |
| *Cs*NikZ | GCGKEQAANNTNEAKGKTLVYGAEMEDEKLNPILIESHAFASDLIFRGLMRFDENNVPKPEIAESYKISDDKLTYDFKIKKGVKFHDNTEVKAEDVVFTVKSILDDKVNSEVKPQFEEIKDVQALNDYEVKVTLKKPFPALLDKMTVGLVPKHALEGKDINTAEFNQKPIGAGPYKFVKWDKGSSLTVERFKDYKGGRPDKVGNIDKVVLKFIPDFNVRAMQLKTGELDATLIDPNQVANLEKESKVTLNKIPTADYRGIMFNFKTKPMFKDVNVRKALNYATDKESIVKGILLGYGFAAYSPIQLNKFNDPDVEKYEYNLDKANELLEQAGWKKGADGIREKDGQKLQFTLFARNNDDTRVKIANYVASQGKKAGFDIKVDARDPKAMKIKDTEAFVIGWGSPFDADDNTYKIFHSSQADHGSGNNFGYYSDPKVDEALEKARTTSDENERKKYYADFQKELADNPAFSMDAYITAIYGVNKKVSGYTTKRVLGHHGEGFLWNTEEWKMQ |
| *Mp*NikZ | ETLVYGSHDYTAINPALYEHGEINSLIFAGLTAHDKDNKLVPALAEKWEYDTATHTWTFHLRDGLKFHDGKELTSEDVKFTLEAILDKKNNSEIISNYQDITKITCPDKKTVVIQLGKENVVFADYMTIGILPKHLLNGKKLATAEFNQKPVGAGPYKLTSWDEGQSITLEKFDGYYAGKPHIKRIIFKIVEDSDARLLQLKSGDLDMAQIEPKQAKSLEKDSKDFDIYRMNTADYRAIAYNFAGSKLFKTYPELATILSYGIDREAIIKSVLLGEGQVAYSPIQKNKFNDSNIEKFCYNPEKMEQLLQQDGWAKNKKGIYEKNGVELKFTITAMADDQVRVDMAKMCAEQLKSHGVSVKAEARKELDWEKQDATIIGWGSPFDADDHTYKIFTSDAGDNYTGYANSTVDDVLAKARSTTNEGERQRYYSQFLSYMTQQMPYTYIAYIDADYAVKKDISGITKDTMLGHHGVGIFWNVADWKLEK |
| *Ng*NikZ | GGGTQDGAGSDTLTYALGDEPDVLNPALVDEHLDPVTEMVFRGLTGHDADNKIVPALAESWKISDDERTYTFSLRDRVTWHDGEPFTADDVVFTIEGIRDGDFVTSNKFADVKDVTADDDHTVTIKLSEPAPALLDTLANGILPEHILAKTGIDDPDFNEHPIGTGPFKLDTWKHDEYATLSAFDDYYDGAPGLDGITISYVPDAATRLIKLKNGEVDAAYLEPQQAKEFDGDGGDGGDNGTEVKVWPTADYRAVMFNMTDDRFKDPAIRQAMNYAVDRDAIIKSVQHGYGSPATGPLDKSPFHADVADYTFDPKRVEKIMTDAGYEKNDDGIWAKDDKPVEFDLTTFAEDGLRASMIEVLATQLREQGFDVTAKPEPQDSVKWDKLETFLIGWGTPYDPDGALYGPFHSSESLAKGGSNYGSYANDDADKALEEGRSTSDEKTRKKAYTDFQKALVEDPPFVFVSYLEAVNAVPSNLEGLQERTLAHHGYGFFWNVEDWSFAK |
| *Su*NikZ | EDTKELTPSNENRLIYASEEEFEGLNPILEETNLDALLFRGVMRFDENNKVVTDIADSFEVSEDKLTYTFKLKEDITFHDGNELTADDIVFTIESIMDDQNASFLKSDFEEVASTNALDDYTVEITLKQPFTPLLDKLTVPILPKHAFEGVDMRTADFNSHPIGAGPYMFDKWDRGNSLTLQAYGDFFGTKPSIEKVIFKFIPDSNVRALQLKSGEVDVALLDPVQVENLEREANLTVYDIESADYRGILFNMNLDLWKDVNVRRAFSYATDREQIVKGILKGHGSVAYSPLQNHEFHNDQIEKYTYDLEKANALLDEAGWKENEDGIRSKDGEKLSFTITAPASDAVRVNMANYVAEGFKDVGADVKVAALDWSAITIEDADAFMVGWGSPYDADHHTHILFHSDESSVKGTGYNYGEYSNEKVDELLAKGRTETDPEERKAIYMQFQEVLANDPAFVFIAYENAVYGINKNVSGVKERTLGHHGSGFLWNVEEWKWNDQ |
| *Tf*NikZ | GCGSQGGKEASKTSVSGKTLIYGAESEFEKINPVLDDSPVDTMIFSGLTKFDENNRPVPDLATSWDISSDNLTYTFHLRKDVKWHDGQPFTANDVKFTIDSILNPKNNSLIKEQFEEIKSVDVIDNYTVKISLKKPFPPILDNLTRGIVPIHILSGKDLNSVDFNSGPIGTGPFKFAEWKKGESFTLQANTGYYGGKPKLDKIVIKFIPDMNVRAIQLETGEIDVALLEPQQLARVEKSEKNKVYLIPTADYRAMMYNFRKPLWQDVKVRQAINYAVNRDNILQGVLLGHGKVAYGPLQLNWANNENVEKYGYNPEKAKALLAEAGWKPGPDGILTKNGQKFSFRLTTFNNDPIRVAIVNALSTELKKIGVEAIPDPRARGTFKWPEVDAFLLGWGSPFDPDDHTYKLFHSSQIENGWNLGAYRDAEIDRLLEEARTTLDENKRKELYGKFQKRLAENPPYNFLVYLDAVWVVNKNVTGIKPKTLGHHGAGFLWNIQEWDKNNGQL |
| *Um*NikZ | GCLGDNKEQVDTAEKKQLIYAVESEIKKVNPILDESQEIDTLLFRGLTKPDENNNPQPDLAESWNISEDNLTYTFQLRKDAVWQDGQPVTAKDVAFTFNKIIDPSTNTPISGEFNQLKSIEATEDYEIVITLHHPYPPLLDKLKVGIVPEHILRNENINETDFNQNPIGNGPFKLKEWKSDDTIILERNSSYYGPAPKLEEVIFKVVPDANTRVLQLKTGEIDLALLEPNQLASIHEDDPFTVHEISTADYRAVMYNLRLPLFQDKRVRQAMNLAVKREELVTGVLAGKGEPAYGPLQKSWAGIPQKEWYTYDPEKAQQLLIEAGWVKGADGILTKDGERFEFELVSPIQDKVRVALANVVSEQLKPLGIIAVPKPIDQHAVDYDGEDALIIGWGSEFDPDDHTYRLFHSNEIGDGKYNFGAYQNQTVDELLIKARTATNTEDRKNYYRKFQEELAIDPAYNFLVYLDALYGVNKNVSGISNRTLGHHGFGILWNIEEWDKE |
| *Vb*NikZ | CGTSSNKSANAEKTLTYASPDCKTINPVLNTHDELPDLIFSGLMKHDGNGKPVVDLAEKYEFDKGTLTYTFHLRKNVKWHDGKPFTAADVKFTLDMLRSNDKLEAEVTDSFKDIKEVTVVDDNTVKIQLSKPNAAMLDNLAIGIMPKHLLEGKDIMTSDFNQHPIGTGRYKFVSWDKGQSIIVQKNDEYYGKAPKISKIVFKIIPDENAKAAQIKSGGVDLALINAKDAKAFRDNKDFKVFDFATADFRAIGPNFKNAYWQNPEHQALIPVLGYALDKKAIVDSVLGGQGEPAYSPLQINKEYNDASVDHRDYNPEVFKQKMEELGWKLGSDGIYEKNGEKLSFSVEAREFEEERVDMAKVASAQFKKLGVDMKVNIVPKFNWKEMQCCLIGQAAPFDPDQGTYDFFVTKASANYTAYSNSAVDAALAAARATYDVAQRKAAYNDFQKAWNAQPAFIMLAYLHGNYVANKKLSGLDTHVILGHHARGVFWNVEDWDIE |

Signal peptides have been removed.

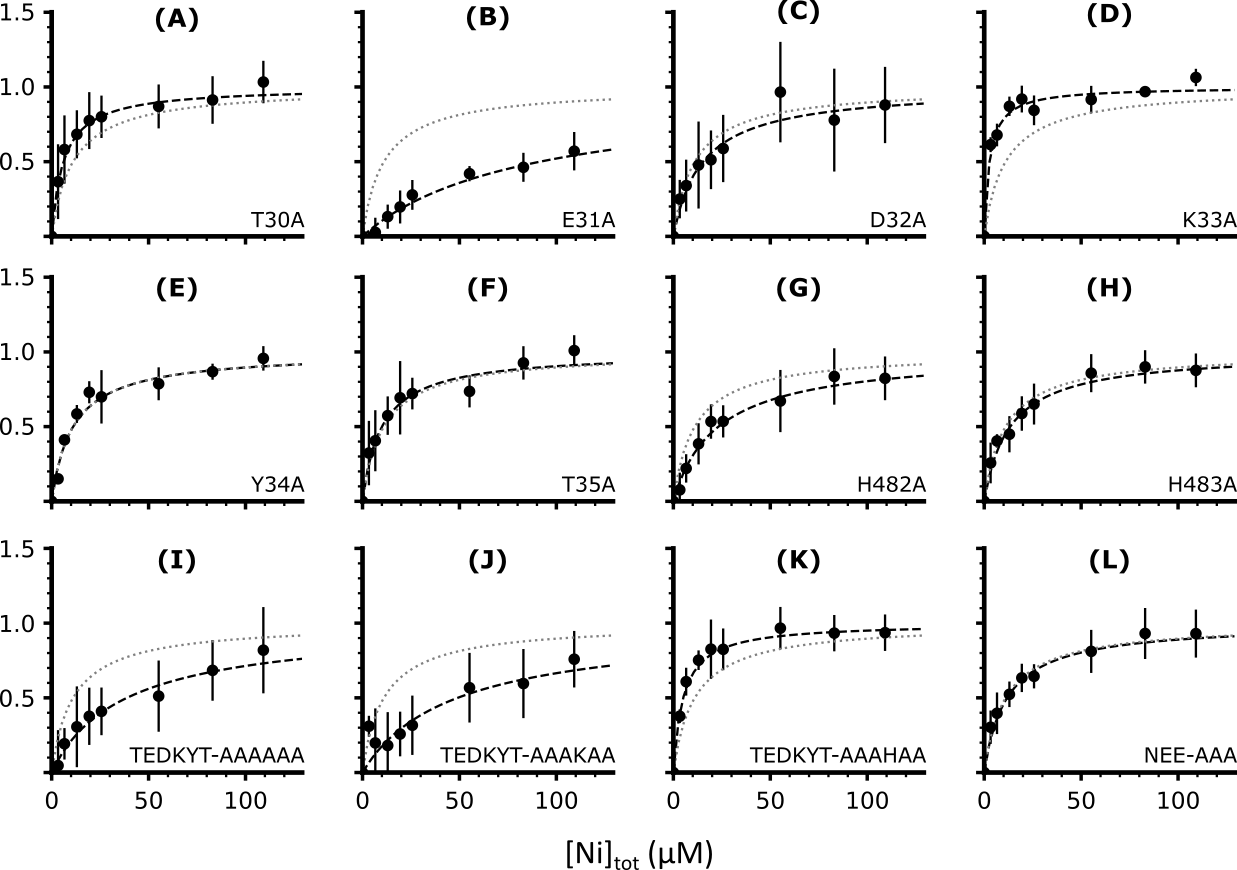

**Figure S1.** **Nickel (Ni^2+^) binding curves of the mutant and engineered *Cc*NikZ-II variants.** The y-axes are normalized to each proteins’ fitted *F*_max­_ value, and all x-axes are the total nickel concentrations throughout the titration in micromolar units. The grey dotted lines represent the curves of best fit for *Cc*NikZ-II to compare the effect of the mutations on the *K*_D_ value.

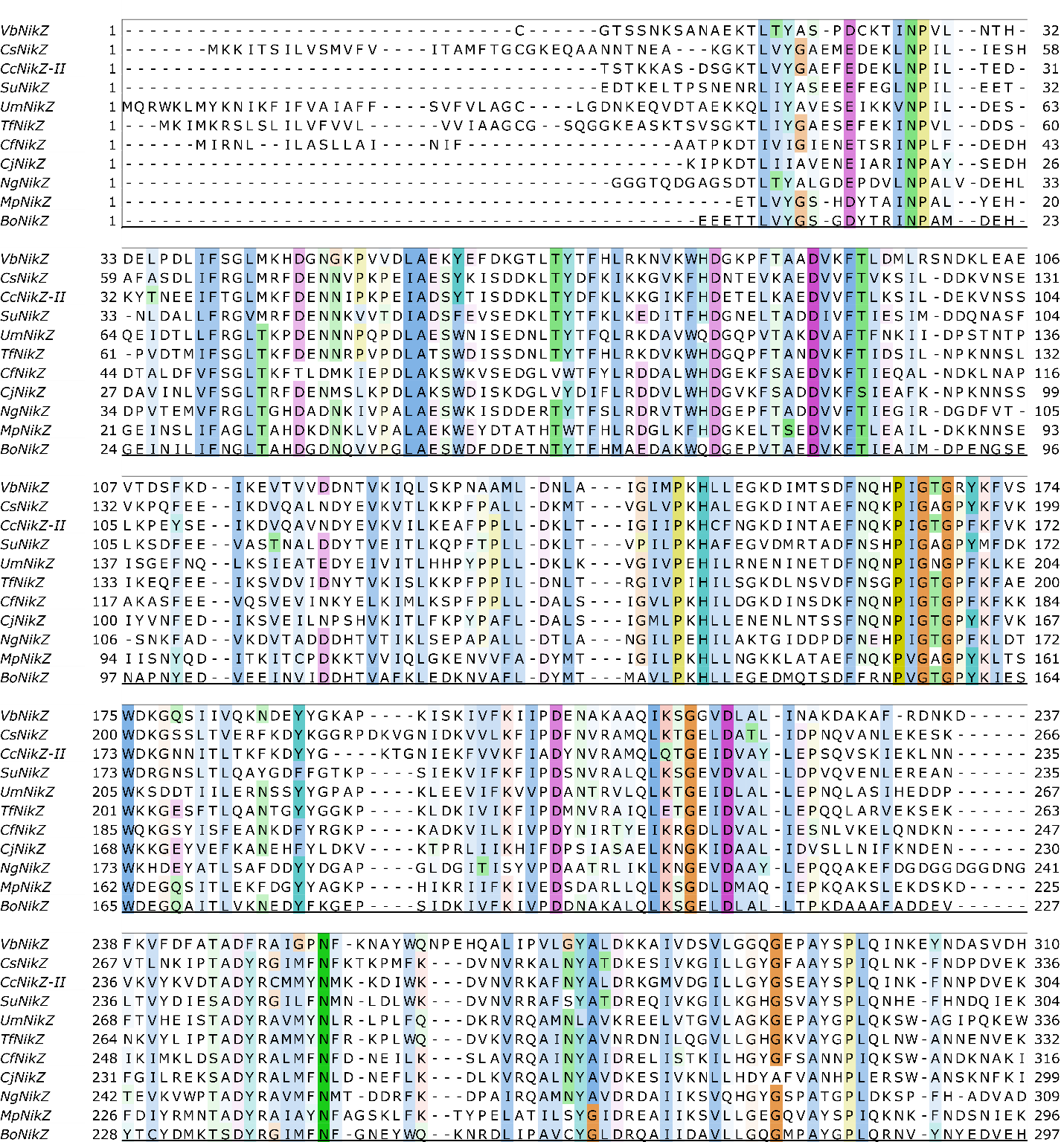

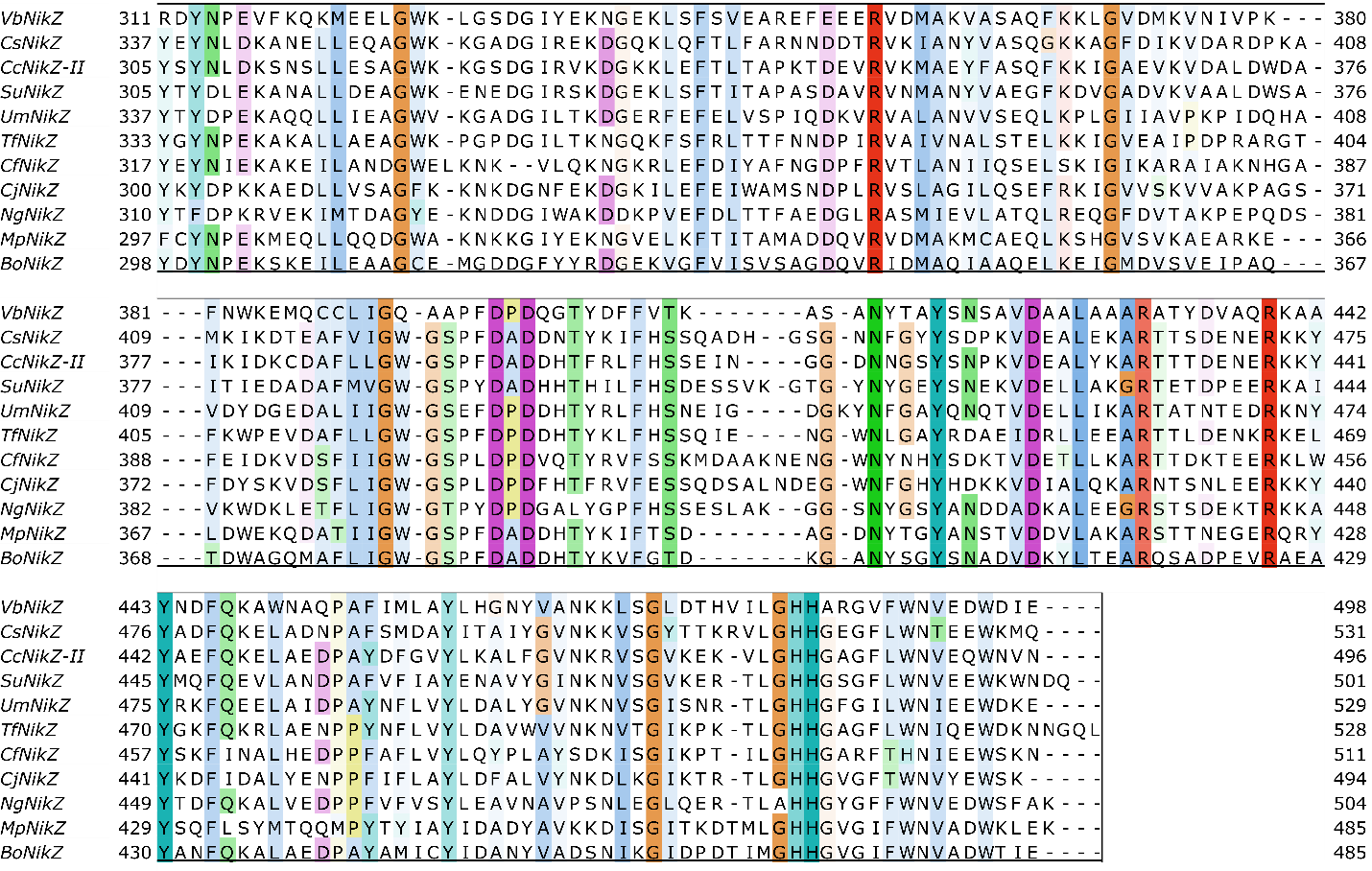

**Figure S2. Full Multiple Sequence Alignment.**

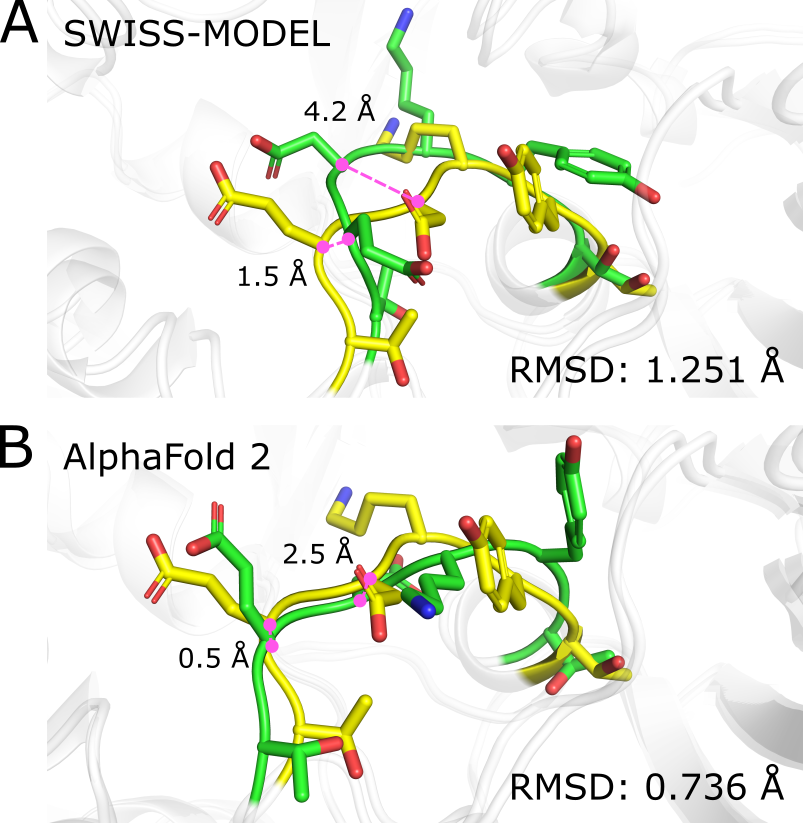

**Figure S3.** Structural alignment of v-loops from the crystal structure of *Cc*NikZ-II and structural models generated using SWISS-MODEL (A) or AlphaFold2 (B). Superimpositions of predicted apo CcNikZ-II (green) to the solved crystal structure (yellow). Magenta dotted line indicates distance between α carbons of aligned residues.

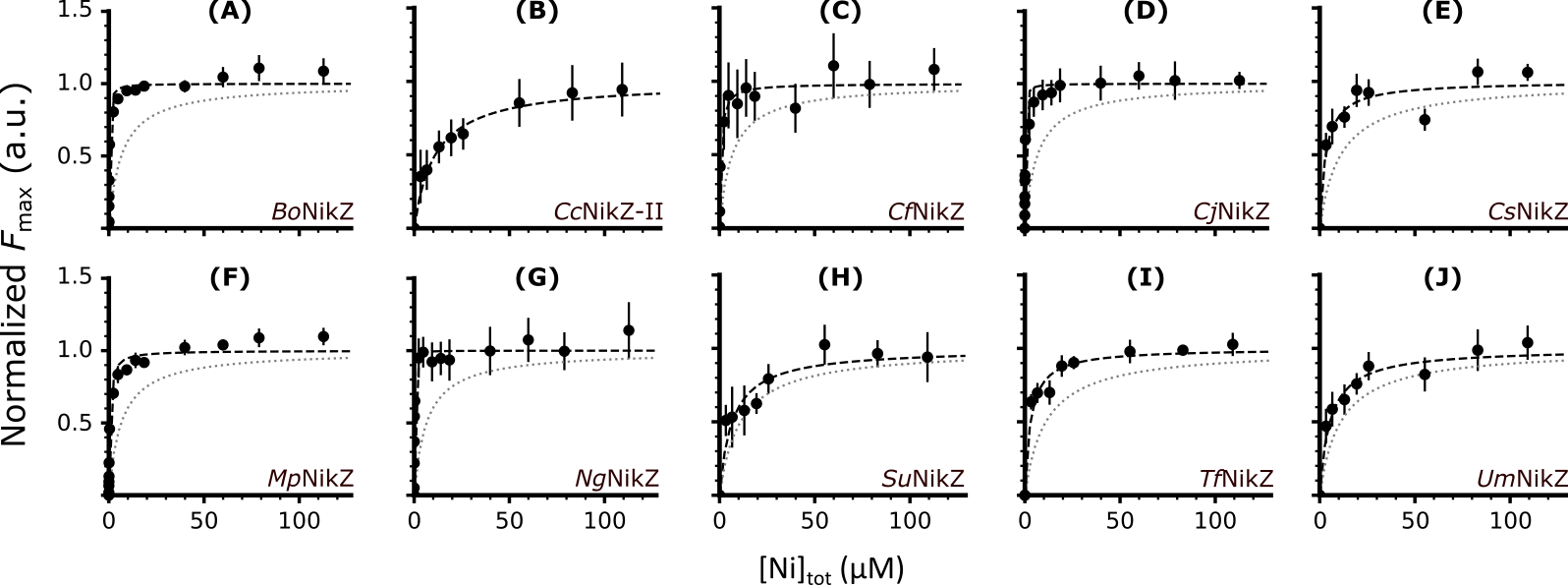

**Figure S4.** Ni^2+^-binding curves of the wild type NiBPs characterized in this study. The y-axes are normalized to each proteins’ fitted *F*_max­_ value, and all x-axes are the total nickel concentrations throughout the titration in micromolar units. The grey dotted line represents the curve of best fit for *Cc*NikZ-II to compare the effect of the mutations on the *K*_D_ value.

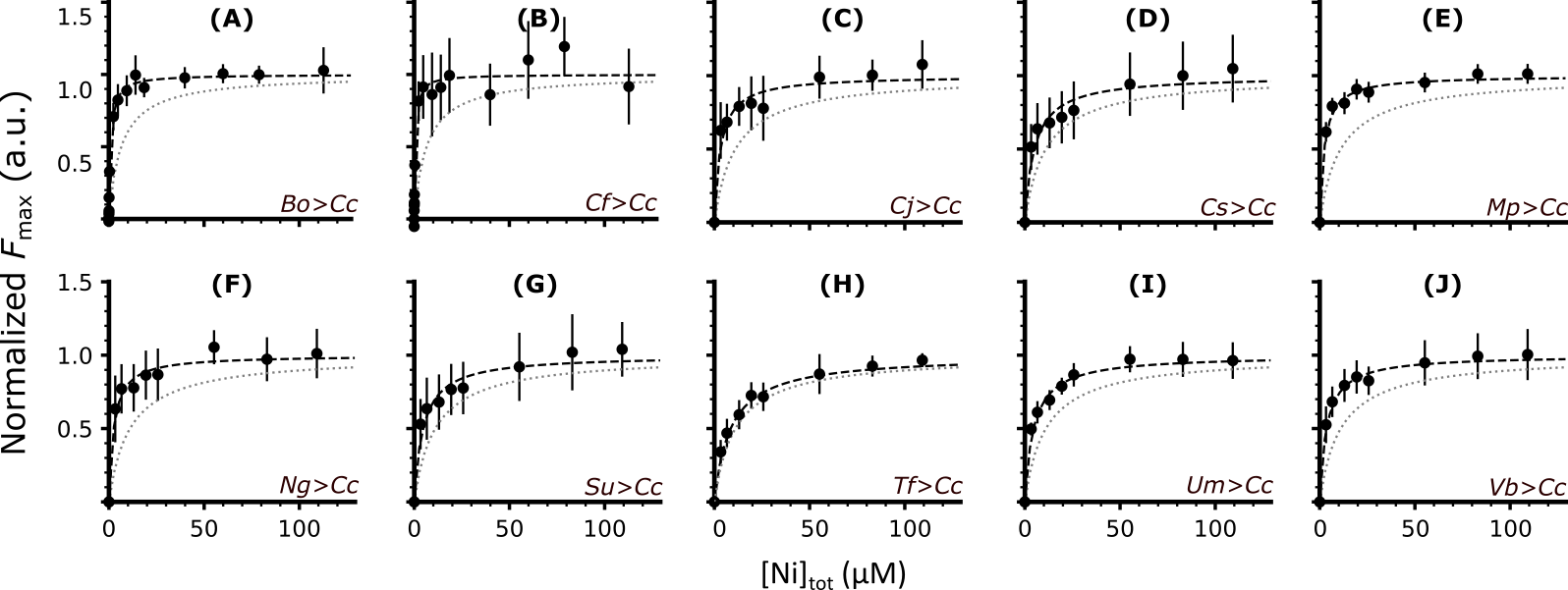

**Figure S5.** Ni^2+^-binding curves of the engineered *Cc*NikZ-II variants with v-loops from other NiBPs**.** The y-axes are normalized to each proteins’ fitted *F*_max­_ value, and all x-axes are the total nickel concentrations throughout the titration in micromolar units. The grey dotted line represents the curve of best fit for *Cc*NikZ-II to compare the effect of the mutations on the *K*_D_ value.

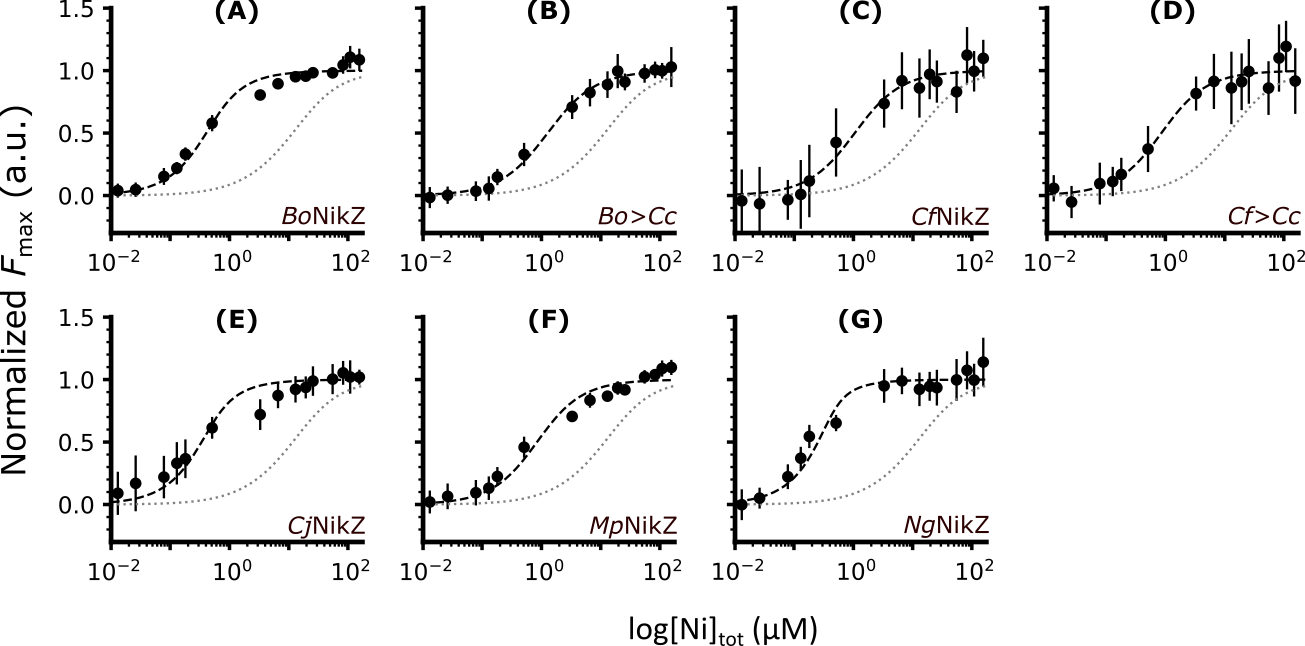

**Figure S6.** Ni^2+^-binding curves of purified wild type NiBPs and engineered *Cc*NikZ-II proteins with the lowest *K*_D_ values (highest affinity) used in this study. The y-axes are normalized to each proteins’ fitted *F*_max­_ value, and all x-axes are the log of the total nickel concentrations throughout the titration in micromolar units. The grey dotted line represents the curve of best fit for *Cc*NikZ-II to compare the effect of the mutations on the *K*_D_ value.

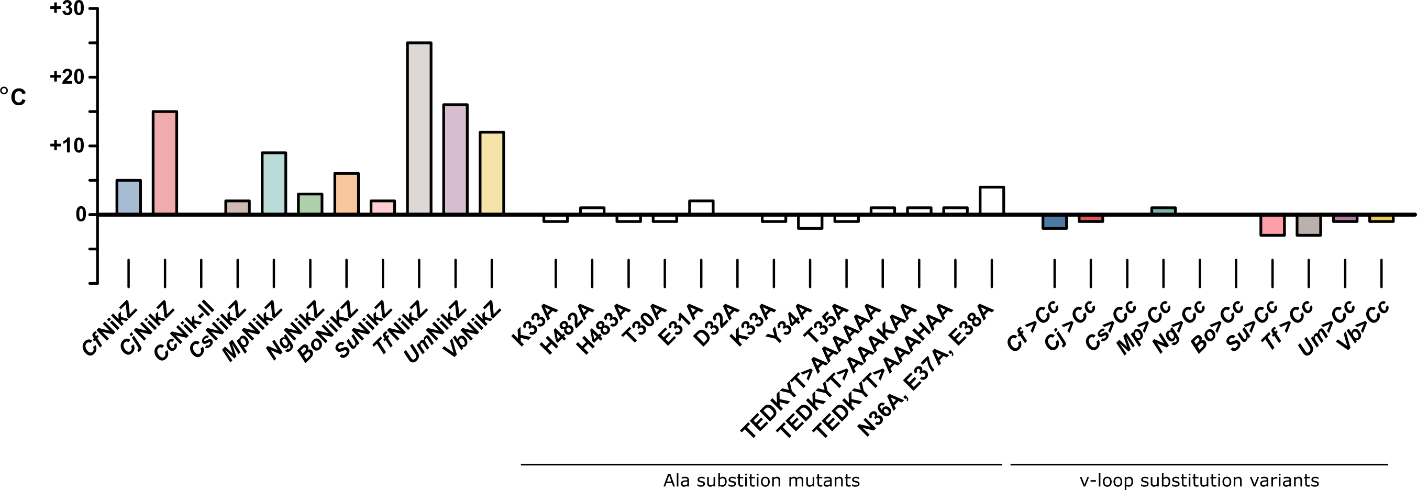

**Figure S7.** Melting points of NiBPs: thermal shift assays with purified wild type and engineered proteins. The *Cc*NikZ-II melting point (48 °C) was subtracted from the melting points of each protein and baselined at 0.

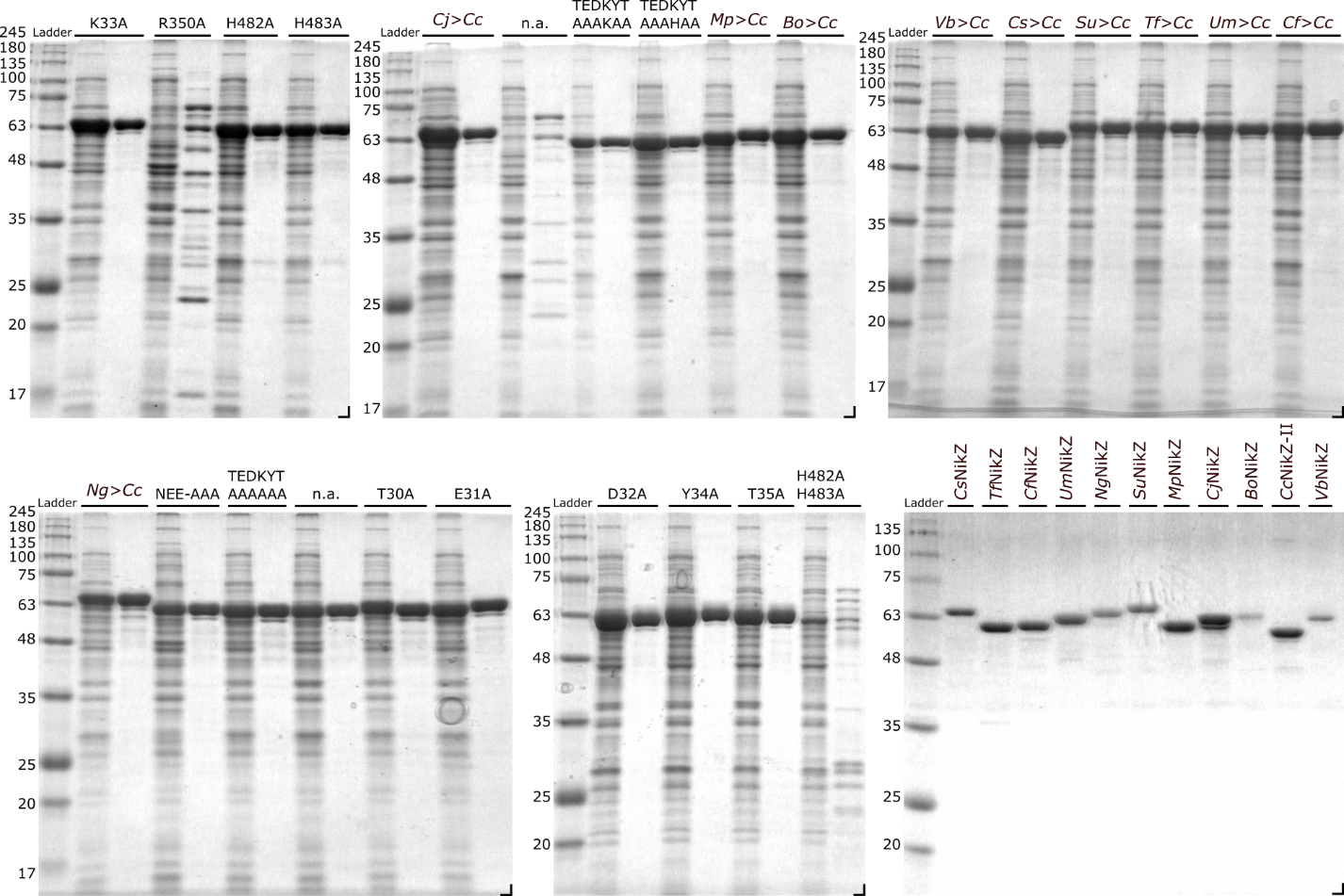

**Figure S8.** SDS-PAGE analysis of purified NiBPs used in this study. Coomassie-stained SDS gels showing purified wild type proteins and engineered *Cc*NikZ-II variants. Ladder, molecular wight markers (kDa). **Additional Supplementary Files**

- Supplementary Data File 1. FASTA file for 4’516 sequences (NiBPs), reduced redundancy
- Supplementary Date File 2. Newick formular for the phylogenetic tree
- Supplementary Data File 3. FASTA file for mutant, homologue, and exchange libraries
